# Predicting selection-response gradients of heat tolerance in a widespread reef-building coral

**DOI:** 10.1101/2021.10.06.463349

**Authors:** Ponchanok Weeriyanun, Rachael B. Collins, Alex Macadam, Hugo Kiff, Janna L. Randle, Kate M. Quigley

## Abstract

Ocean temperatures continue to rise due to climate change but it is unclear if heat tolerance of marine organisms will keep pace with warming. Understanding how tolerance scales from individuals to species and quantifying adaptive potentials is essential to forecasting responses to warming. We reproductively crossed corals from a globally distributed species (*Acropora tenuis*) on the Great Barrier Reef (Australia) from three thermally distinct reefs to create 85 offspring lineages. Individuals were experimentally exposed to temperatures (27.5, 31, and 35.5 °C) in adult and two critical early life stages (larval and settlement) to assess acquired heat tolerance via outcrossing of offspring phenotypes by comparing five physiological responses (photosynthetic yields, bleaching, necrosis, settlement, and survival). Adaptive potentials and physiological reaction norms were calculated across three stages to integrate heat tolerance at different biological scales. Selective breeding improved larval survival to heat by 1.5 - 2.5x but did not result in substantial enhancement of settlement, although population crosses were significantly different. At heat, adults were less variable compared to larval responses in warmer reefs compared to the cooler reef. Adults and offspring also differed in their mean population responses, likely underpinned by heat stress imposing strong divergent selection on adults. These results have implications for downstream selection during reproduction, evidenced by variability in a conserved heat tolerance response across offspring lineages. These results inform our ability to forecast the impacts of climate change on wild populations of corals and will aid in developing novel conservation tools like the assisted evolution of at-risk species.

**SUMMARY STATEMENT:** Heat stress exerts disruptive selection on adult corals. This likely underpins variability in offspring survival and results in differences in offspring responses to selection.

## INTRODUCTION

Ecosystems globally continue to suffer from the adverse effects of anthropogenic climate change. Coral reef ecosystems in particular are declining as sea surface temperatures (SSTs) have risen dramatically (Hughes et al., 2017). Coral reefs are some of the most productive habitats and the continued loss of corals will likely impact reef community dynamics by changing ecological functioning via trophic interactions, with downstream impacts on fish and other invertebrate species (Jones et al., 2004; Komyakova et al., 2013; Nelson et al., 2016). Coral reefs are biodiversity hotspots that provide access to food, act as natural barriers against storm damage and benefit the economy through tourism and other recreational activities (O’Mahony et al., 2017). Coral reef ecosystems like the Great Barrier Reef (GBR) contribute, on average, $6.4 billion (AUD) to the Australian economy per annum (O’Mahony et al., 2017). Consequently, the loss of these crucial habitats will have devastating economic and social-ecological effects on coastal communities.

The upper thermal thresholds of reef-building corals generally reside ~1 °C higher than their local summer maxima (Schoepf et al., 2019). Once temperatures exceed this threshold, corals may lose their symbiotic dinoflagellates (Symbiodiniaceae) through a process known as bleaching, which is typically experienced by corals during marine heatwaves if warming persists for extended periods of time (Hoegh-Guldberg, 1999). Extreme bleaching (defined as >60 % of bleached corals present within a reef) often leads to high mortality (Hughes et al., 2017). At least four mass bleaching events have been recorded on the GBR in the last century - 1998, 2002, 2016 and 2017 – generally increasing in intensity and severity (Hughes et al., 2017) and have led to large-scale losses in hard coral cover. Rates of decline vary by region, but overall, declines have rapidly increased per year since 2006 (De’Ath et al., 2012), although there have been recent signs of recovery. Therefore, bleaching-related stress has not been equal across the GBR. This trend in decline is reflected globally, with SSTs continuing to exceed previously held records (Heron et al., 2016), suggesting that corals may soon reach their physiological limits to cope with increasing heat (Matz et al., 2017).

Changes in temperature have the potential to impact organismal life-stages differently, in which larvae, recruitment stages, and adults can all vary in their thresholds to climate extremes. Despite this, information on larval physiological responses is relatively unknown for coral early life-history stages (McLachlan et al., 2020), with 95 % of studies focusing on adult responses and only 2 % and 1 % on pre-settled and settlement stages, respectively (McLachlan et al., 2020). In corals, sexually mature adults release their gametes within a temperature range of ~28 – 30 °C, with larvae of some coral species able to survive 2 – 5 °C above this range (Heyward and Negri, 2010). However, coral spawning typically occurs during the warmest summer months, when spikes in SSTs are most likely to occur (Keith et al., 2016). As a result, coral larvae are potentially subjected to much greater temperatures, increasing their risk of mortality and reducing recruitment and settlement success (Heyward and Negri, 2010). After fertilisation, larvae undergo a pre-competency period where they develop and disperse throughout the water column. Once competent, they respond to biophysical and chemical cues to find optimal settling conditions (Doropoulos et al., 2018). Increased SSTs may induce premature metamorphosis by increasing larval metabolic activity and development rates, thereby reducing pre-competency periods (Heyward and Negri, 2010). This reduction may influence subsequent larval dispersal distances and result in settlement occurring in suboptimal conditions (Edmunds et al., 2001). Therefore, it is important to incorporate these responses into predictive models of ecosystem change given their flow-on effects into key demographic and population level dynamics.

Reductions in dispersal distances have the potential to reduce gene flow between populations, thereby potentially accelerating population differentiation and local adaption (Bassim et al., 2002). Additionally, some coral populations rely on the dispersal of larvae from neighbouring populations for the addition of beneficial genes/genotypes to their gene pool (Munday et al., 2009; Quigley et al. 2019). With reduced reef connectivity, populations may not be able to adapt rapidly enough to cope with increasing SSTs (Quigley et al. 2019) or may struggle to regenerate following mass bleaching and mortality (Bassim et al., 2002). There is scope for adaptation of corals to increasing temperatures (Matz et al., 2018). Further, the temperatures at which bleaching is occurring have increased by ~0.5 °C from 1998 to 2017, suggesting an increase in more heat-adapted genotypes within coral populations (Sully et al., 2019), potentially through processes like selective sweeps (Quigley et al., 2019a). Locally, there have been reported increases in coral cover in both the central and southern GBR (AIMS, 2020), with evidence to suggest that warm or “extreme” habitats harbour an increased number of thermally adapted genotypes with the potential to transmit heat tolerance (Quigley et al., 2020a; Schoepf et al., 2019). However, current estimates of the rate of SST increases suggest that these increases may exceed the potential rate of fixation of beneficial genetic variants given factors like currents and reef topology (Quigley et al., 2019a). Additionally, the annual increase in SSTs has extended the period at which ‘summer’ temperatures occur, further increasing global bleaching by reducing potential recovery periods that occur when coral populations are exposed to cooler ‘winter’ temperatures (Heron et al., 2016). Taken together, this suggests that corals adaptive potentials may be constrained.

Variability in heat tolerance has been observed at many ecological levels, including between individuals, populations, species, and by reef region. For example, differences in the heat tolerance of corals have been observed across reefs globally, with coral populations present along the Persian Gulf having demonstrated very high thermal thresholds of ~ 4 °C above mean monthly temperatures (Kirk et al., 2018; Moghaddam et al., 2021; Savary et al., 2021). Northern GBR corals also demonstrate higher thermal tolerances compared to some central populations, in which heritable host genetic mechanisms play a role in the thermal resistance of these northern corals (Dixon et al., 2015; Quigley et al., 2020a). This geographically distinct habitat could contribute to sub-speciation, seen in other marine invertebrates that have different thermal thresholds, like subspecies of *Crassostrea gigas* (Ghaffari et al., 2019). However, it is less clear whether similar patterns in heat tolerance at the individual coral genotype scale to the population level, a trend seen in fish, but not insect or plant species collected across temperature clines (Payne et al., 2021; Rezende and Bozinovic, 2019). Quantifying the scaling of heat tolerance across different biological levels will therefore help to elucidate the baseline potential of wild populations to adapt. Further, evolutionary models incorporating metrics like narrow-sense heritability (h^2^; the phenotypic traits within an organism that arise from allele inheritance; Evans et al., 2018), selection coefficients (S; relative fitness of a phenotypic trait; van Tienderen and de Jong, 1994), and responses to selection are commonly measured (Falconer and Makcay, 1996), and allow for the prediction of organismal responses to future stressors. Thus far, models incorporating the “breeders equation” (R = h^2^S; used to predict the effect of selection pressures on phenotypic traits; Falconer and Makcay, 1996) have demonstrated the utility of evolutionary modelling for predicting the potential of the coral’s algal symbionts to confer increased survival (Quigley et al., 2018). Similarly, these principles can be applied to the coral host to quantify adaptive potentials, in which this information can then be used to harness the existing adaptive potential of coral populations for conservation and restoration.

Coral restoration interventions, some of which can be classified as assisted evolution methods (van Oppen et al., 2015), aim to increase the adaptive potential of organisms by targeting the acceleration of adaptation through various genetic mechanisms. This may include selective breeding for the selection of desirable phenotypes (van Oppen et al., 2015). Assisted Gene Flow (AGF) is one intervention that utilises the differing thermal tolerances of parental colonies combined with the intentional movement of recombinant offspring, by reproductively mixing populations sourced from different environmental conditions within the same species to prepare future populations for increased warming. This strategy is based on increasing gene flow between populations that have a beneficial phenotype(s) with those of a target population(s) via the creation and subsequent introduction of new, more resilient genotypes (Aitken and Whitlock, 2013). Selective breeding through the outcrossing of gametes between these reefs of distinct thermal profiles assumes that populations exposed to climates of similar temperatures to those predicted to arise from anthropogenic climate change may be better adapted, and therefore better able to cope with warming (Aitken and Whitlock, 2013). The inheritance of these beneficial alleles by populations that evolutionarily have experienced cooler temperatures should increase the overall resilience of the species via increased population persistence (National Academies of Science, Engineering and Medicine, 2019).

Here, we compare the physiological responses at heat stress of selectively bred *Acropora tenuis* larvae and newly settled recruits and their parental colonies collected from three sites along the Great Barrier Reef – Davies and Esk reefs (central GBR) and in the Keppels (southern GBR). Percentage larval survival and settlement were recorded following exposure to control (27 °C) and heat stress conditions (35.5 - 36 °C) and compared to adult heat tolerance at 31 °C, as measured in bleaching score, photosynthetic quantum yields, percent necrosis, and percent survival. We then compared these responses across multiple biological scales (individuals to populations) to calculate adaptive potentials from data collected across each life stage. This work contributes to our understanding of the potential to increase heat tolerance and for the transgenerational transfer of this tolerance using the selective breeding of coral populations sourced from along the GBR.

## MATERIALS AND METHODS

### Coral colony collections

Reproductively mature *Acropora tenuis* colonies were collected in early November 2019 from three sites on the GBR in Australia, including two central reefs: Davies (−18.90, 147.87) and Esk (−18.86, 146.62), and one southern reef in the Keppel islands (23.06, 150.95). Corals were identified as individual genotypes during collection by removing colonies approximately 10 m apart if possible. Genotypes were not known *a priori*. Esk was on average the warmest reef location, followed by Davies, and then the Keppels (**Fig. 1A**). The mean maximum monthly temperature of each site was as follows: Esk 26.26 ±2.30 °C, Davies 26.06 ±1.80 °C, and the Keppels 24.62 ±2.30 °C (**Fig. 1B**). Corals were transported by boat to the National Sea Simulator at the Australian Institute of Marine Science (AIMS). Colonies from each location were acclimated in separate outdoor holding tanks under constant 0.2 μm filtered seawater (FSW) flow-through conditions, maintained at 27.5 °C.

**Figure 1.**
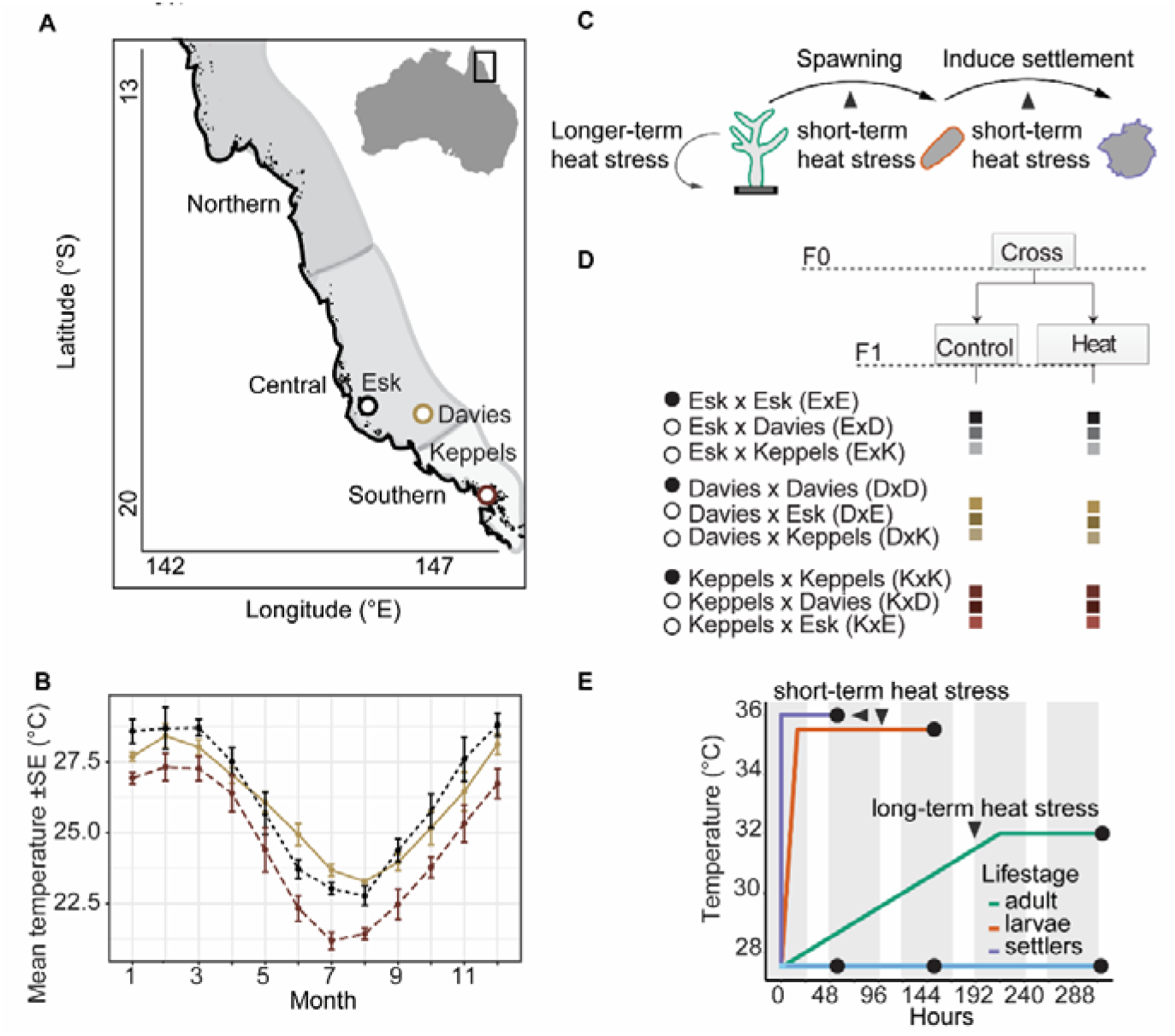
Summary of experimental design for determining variability in heat tolerance across coral life-history stages. Map of the Great Barrier Reef showing the three sites of adult coral collection (A), including Esk (central inshore, black), Davies (central mid-shelf, tan) and Keppels (southern inshore, maroon) reefs (A). Mean monthly sea surface temperature profiles at each reef of adult coral collection (B). A graphic showing when the three life-stages were experimentally exposed to heat stress (C). Summary of the population crosses produced from the three coral populations. Filled circles represent intrapopulation crosses, whereas open circles represent interpopulation crosses (D). Figure indicating the ramping time to reach each experimental temperature and the length that it was held (E). Circles indicate the end of each experimental time point, triangles indicate specific points along the experimental timeframe.

### Coral spawning, selective breeding, and larval rearing

Prior to spawning, each *A. tenuis* colony was given an identification number (**Table S1)**. Individual colonies were isolated in separate bins following signs of spawning imminence, which includes the appearance of egg–sperm bundles under the oral disc and “setting” (polyps extended but tentacles retracted). Gamete bundles were released between 19:00 and 19:40 on the 17^th^ and 18^th^ of November 2019 and collected by gently skimming and collecting the bundles off the surface of the water. The eggs and sperm in the bundles were separated by washing for ~2 mins with FSW through a 120 μm mesh filter. The number of sperm was quantified using a Computer Assisted Semen Analyser (CASA) (software CEROS II from Hamilton Thorne). The sperm from one parental colony was then added at a concentration of 10^5^ mL to isolated and cleaned eggs from another parental colony to create 85 distinct coral families with at least one cross per family. These families, observed as biological replicates, comprising of intrapopulation (within the same reef) and interpopulation (between different reefs) crosses, where both are generally referred to here as population crosses (**Table S2**). Population-level crosses are referred to as follows with the maternal colony first and then paternal colony, in which intrapopulation crosses (filled circles; **Fig. 1D**) are ExE (EskxEsk), KxK (KeppelsxKeppels), DxD (DaviesxDavies), and interpopulation crosses are ExK (EskxKeppels), ExD (EskxDavies), KxE (KeppelsxEsk), KxD (KeppelsxDavies), DxE (DaviesxEsk), and DxK (DaviesxKeppels) (open circles; **Fig. 1D**).

Eggs were allowed to fertilize in separate bowls per cross for 3 hours. Every hour aliquots of developing embryos were taken from each family (hereafter referred to as crosses; **Fig. 1C-D**) to visually inspect fertilization success and cell division under magnification. Once fertilization was confirmed, embryos from each separate bowl were transferred into separate 15 L constant flow-through conical tanks with 0.2 μm FSW in a temperature-controlled room, such that the temperature of each cone was maintained at 27.5 °C, *p*CO_2_ 400±60 ppm, ambient light (i.e. non-photosynthetic), and salinity of 35 psu. Each flow-through cross culture had an outflow covered by a 10 μm filter and air curtain of bubbles to prevent larvae from collecting on the outflow filter. By days three and four post-fertilisation, larvae were ciliated and motile, consistent with the 96-hour stage of larval development.

### Larval heat stress experiment

Thirty larvae from each of the total 85 crosses were placed into floating net-wells (n = 3 replicates per cross, per temperature), separated into two holding tanks. One tank was set at 27.5 °C (control treatment) and the other at 35.5 °C (heat treatment). To achieve this temperature, the heat treatment was ramped up to 35.5 °C in hourly increments of 0.5 °C from 27.5 °C. Once 35.5 °C was reached, survival assessments began, in which the number of larvae that were alive in each net-well (of the total n = 30) was counted twice daily to estimate survival rates per replicate. The final survival counts occurred once ~50 % of the larval crosses reached 50 % survival in the heat treatment. No crosses perished post-fertilization before sampling took place.

### Settlement experiment

Following the larval heat stress experiment, only 69 of the original 85 crosses contained sufficient larval stock for further experimental use. These 69 crosses were used to investigate the effect of thermal stress on larval settlement behaviour. Larvae were sampled 35 days after spawning and transferred to sterile six-well plates (n = 10 larvae per well, n = 3 wells per cross, n = 2 crosses per sterile six-well plate) containing 10 mL of 0.2 μm FSW. Crustose coralline algae (CCA) rubble was freshly cut into 3 × 3 mm chips using bone cutters, and placed into each well to induce larval settlement. The plates were then placed into plastic bags and sealed to prevent evaporation and replicates were transferred into two incubators with photosynthetic lights (12:12 day:night light cycle, 170 to 180 PAR, Steridium E-500, lights: Sylvania FHO24W/T5/865 and Innova 4230, light: Sylvania F15W/865) set at 27 - 27.5 °C (control treatment) and 35.5 - 36°C (heat treatment). The number of settled larvae (metamorphosed and attached to substrate) were counted for 17, 24 and 48 hours after incubation at both temperatures and the number of settled larvae was recorded. Settled larvae were defined as those that were attached to the well or CCA and deposited a basal plate that was visible after metamorphosis. This was distinguished from only metamorphosed larvae (metamorphosis but no attachment to substrate), which were excluded from this analysis.

### Adult heat stress experiment

Colonies were kept in outdoor aquaria under the following conditions before fragmentation: 0.2 μm FSW, 27.5 °C, *p*CO_2_ 400±60 ppm, and salinity of 35 psu. Three to five colonies representing different genotypes for each population were fragmented using a “diamond-tipped” bandsaw (**Table S1**). Each colony was divided into a minimum of six fragments, with each placed into one of six experimental tanks (n = 3 control tanks, n = 3 heat tanks). For the Davies population, five genotypes were selected and cut into 82 fragments; three genotypes were used from the Esk population (51 fragments); and three genotypes from Keppels population (28 fragments). Each fragment was glued to a calcium carbonate plug using epoxy glue, placed into PVC plug holders (hereafter referred to as “sticks”), and allowed to acclimate at control temperatures before being moved to the experimental tanks. This acclimation period attempts to minimize the effects of sampling and handling before the experiment, where all coral fragments were allowed to acclimate and recover from the fragmentation stress for one week. During this time and over the course of the experiments, corals were fed once a day at 16:30 hours with 0.5 individual/mL of *Artemia* sp. and 5×10^6^ cells/mL of microalgae (*Parmelia sulcate*, T-ISO, *Chaetoceros muelleri, Nannochloropsis oceania*, and *Dunnaiella sp*.).

Upon moving the corals to experimental tanks, the position of each fragment was randomized in each tank and across all tank replicates, such that each tank included all genotypes from each population. Experimental tanks were filled with 45 L of either 27.5 °C FSW (control treatment) or 31 °C (heat treatment) on constant flow through. Each tank was equipped with a tungsten aquarium pump for constant aeration and mixture of water. The heat treatment corals were ramped at a rate of 0.5 °C until 31 °C was reached. The light cycle for each tank was set on a 12:12 hour regime and light intensity at 171 PAR. Sunrise was set at 09:00 h. The coral fragments underwent these experimental conditions for 14 days. Temperatures for larval and adult experiments were chosen in order to compare with previous work performed in the same region using a cross design (Dixon et al. 2015; Quigley et al. 2020b).

Coral colour, as a proxy for bleaching, was determined using an underwater colour reference card for corals (Coral Watch Card; Siebeck et al., 2008). Images with the bleaching card were taken at least twice a week, and fragments were compared to the brown hue (D1-D6). ImageJ (Schneider et al., 2012) was used to create a Red-Green-Blue standard curve for coral colour as per Quigley et al. (2019b). Survival of each coral fragment was noted from the images. Coral death was determined by the presence of microalgae growing on the bare skeleton on each coral fragment. The percentage of tissue necrosis was measured using the ImageJ surface area tool (Schneider et al., 2012) per coral fragment, per tank, per time point. Effective quantum yield of photosystem II (ΔF/FM’), which is the efficiency of photon absorption, was measured using the Diving PAM (Walz, 2018). PAM measurements of each fragment were taken twice per week at 10:00 h.

### Statistical analysis

#### Larval heat stress experiment

After testing data normality and homogeneity of variance with diagnostic plot using “stats” package (R Core Team, 2021), differences in larval survival were assessed using the non-parametric *“wilcox*.*test”* from the package “ggpubr” (Kassambaraype, 2020) to compare if the mean values between each cross at the control and heat treatments (two independent groups) were statistically different.

#### Settlement experiment

The percentage of settled larvae was first analyzed using the base “stats” package (R Core Team, 2021) to test the normality and homogeneity of the values. If non-normal distributions were present, as shown through diagnostic plots, the random factors “cross” and “plate” were tested for their contribution to variability in settlement. The “ggpubr” package (Kassambaraype, 2020) and the Wilcoxon’s test (Wilcoxon, 1945) were used to statistically compare the median percentage settlement between heat and control treatments at each time point.

#### Adult heat stress experiment

Several coral traits have been identified as important biometrics for restoration, including partial mortality and bleaching, in conjunction with classic response measurements like survival (Baums et al., 2019). Here bleaching was assessed by the change in colour from photographs, where a “6” is indicative of a non-bleached, healthy fragment and a “0” is a white, bleached fragment. Differences in the photophysiological responses of the algal symbionts within corals (ΔF/FM’), percent necrosis, bleaching and survival were evaluated with respect to temperature treatment and coral population using linear models implemented in the “lme4” package (Bates, 2005).

The metric ΔF/FM’ and the percentage of necrosis were treated as continuous variables, and temperature treatment and population were set as fixed factors in each model. Replicate tank and fragment holder (“sticks”) were set as random effects. These factors were not significant and were dropped from the final model (**Table S3**). Once these random effects were dropped, the linear model was re-fit, and all model assumptions were checked (linearity, normality, and homogeneity) using diagnostic plots in the “stats” package (R Core Team, 2021). Finally, the negative binomial generalized linear model was used for bleaching, linear model for ΔF/FM’ and the percentage of necrosis, and generalized linear model for survival. The function “*anova”* was used to calculate model p-values, used for interpreting the significant difference among means. Post-hoc pairwise comparisons of population and temperature treatment were then run on the model outputs (Tukey, 1977). For survival data, a binomial distribution was used to see if there was a significant difference between populations and temperature treatments. For each trait, the statistical difference in median values was assessed using the packages “ggplot2”, and “plyr” (Wickham, 2011; Wickham, 2016), with the Wilcoxon’s test (Wilcoxon, 1945). The percentage change of adult responses by latitude was calculated and plotted using “ggplot2” package (Wickham, 2016). All analyses were carried out using RStudio, version 2.13.2 (R Core Team, 2021).

#### Predicting response gradients to selection

The density plots of adult and larval survival were made by using the “ggplot2” package (Wickham, 2016). The breeder’s equation, R = h^2^S (Falconer and Makcay, 1996) was used to calculate expected responses to selection (R) and potential constraints to the evolution of heat tolerance given selective potentials (S) and narrow-sense heritability for this trait (h^2^). This approach has previously been used in corals (Quigley et al., 2018) to determine how selection on host-symbiont communities may adapt to different selective environments. Contour plots of R, S and h^2^ for larvae grouped by maternal reef of origin were made using the packages “plotly” (Plotly Technologies Inc., 2015) and “dplyr” (Hadley Wickham et al., 2020).

## RESULTS

### Differences in larval survival under heat stress

The influence of both temperature treatments on larval survival varied by population cross, where larval lineages produced from Davies, Esk and Keppels corals displayed both high and low survival under control and heat conditions (**Fig. 2A**). Both median survival and the variance around the median at both control and heat conditions varied across the lineages. Overall, the Keppels intrapopulation larvae survived better at heat compared to interpopulation larvae, whilst the Esk and Davies larvae survived better when crossed with either of the other two reefs as interpopulation crosses. The overall “winners” at heat were DaviesxEsk and EskxKeppels (which also had high survival at control), while the overall losers at heat were KeppelsxDavies and KeppelsxEsk.

**Figure 2.**
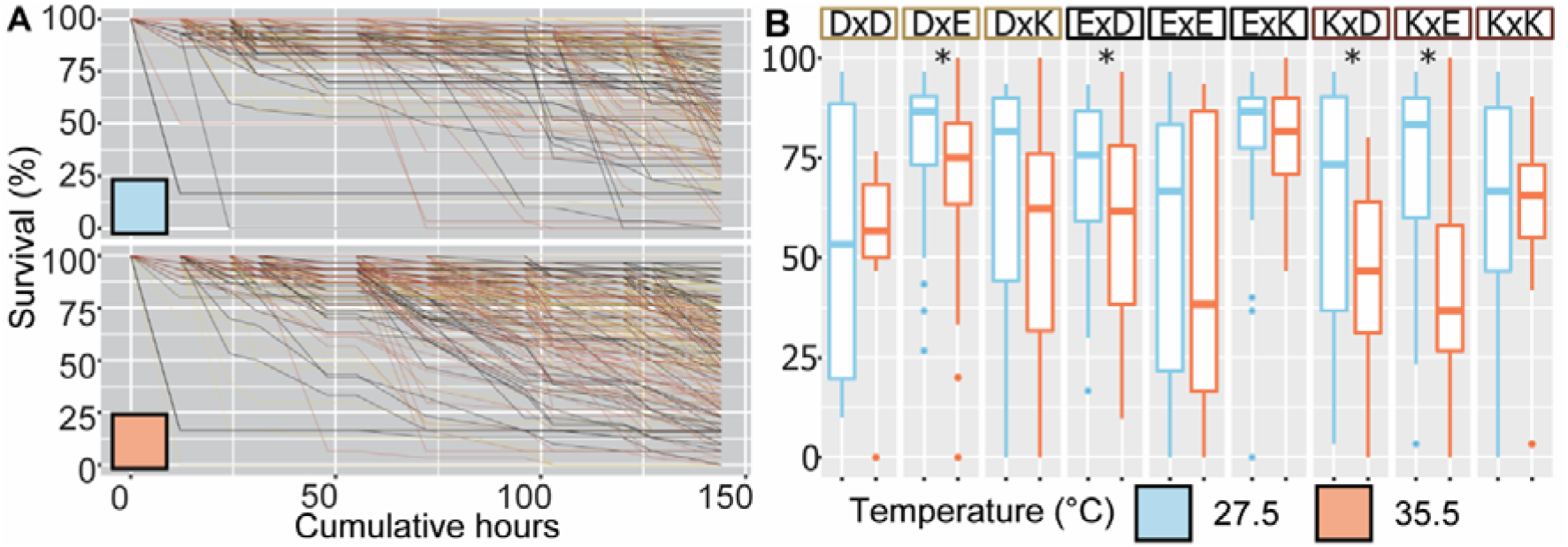
Larval survival under experimental stress. Percentage survival over time of n =85 different *Acropora tenuis* lineages of larvae exposed to control (27.5 ºC) and heat stress (35.5 ºC) (A). The coloured boxes around the crosses indicate the source reef of the maternal colonies: Davies (tan), Esk (black), and Keppels (maroon) (B). The Wilcoxon’s test was performed to analyze statistical differences.

When scaled to the population level, Davies purebred larvae and larvae produced using Davies eggs were only significantly different in their survival between temperatures for one of the three lineages (**Fig. 2B**, *P =* 0.71, 2.50×10^−2^, 9.70×10^−2^ for DaviesxDavies, DaviesxEsk, and DaviesxKeppels, respectively). Survival at heat was higher in the interpopulation crosses compared to the purebred larvae (median survival: DaviesxDavies = 56.70 %, DaviesxEsk = 75.00 %, DaviesxKeppels = 62.30 %). Again, Esk purebred larvae and larvae produced with Esk eggs were only significantly different in their survival between temperatures for one of the three lineages, with the same (reciprocal) cross significantly different (*P =* 4.10×10^−2^, 0.53, 0.29 for EskxDavies, EskxEsk, and EskxKeppels, respectively). Survival at heat was higher for interpopulation crosses than the purebred Esk larvae (EskxDavies = 61.70 %, EskxEsk = 38.30 %, EskxKeppels = 81.70 %). The variation in median survival was also greater in purebred Esk larvae compared to other lineages at heat. Keppels interpopulation larvae all survived less at heat compared to control conditions (*P =* 3.20×10^−2^ and 2.30×10^−8^ for KeppelsxDavies and KeppelsxEsk) whilst purebred larvae survived about equally between treatments (*P =* 0.61). Median survival was lower in the interpopulation larvae compared to purebred larvae (KeppelsxKeppels= 65.60 %, KeppelsxDavies = 46.70 %, KeppelsxEsk = 36.70 %).

### Influence of heat stress on larval settlement rates

When exposed to heat, larvae from all crosses significantly decreased in their settlement behaviour (attachment and metamorphosis) relative to the control temperature after 17 (F_1,433_= 370.30, *P* = 2.00×10^−16^), 24 (6/9 crosses with 0.00 % settlement; F_1,430_= 916.50, *P* = 2.00×10^−16^), and 48 hours (7/9 crosses with 0.00 % settlement; F_1,334_= 722.80, *P* = 2.00×10^−16^; **Fig. 3A-C**). Specifically, larvae from all crosses settled significantly less at heat compared to control temperatures, regardless of population cross (*P*-values in **Table S4**). After 48 hours, DaviesxEsk and EskxKeppels showed the greatest percentage settlement compared to the other crosses (median settlement at heat = both 0.00 %, control = 60.00 and 75.00 %, respectively), in which DaviesxEsk was the only population cross that settled at heat (upper quartile = 100%).

**Figure 3.**
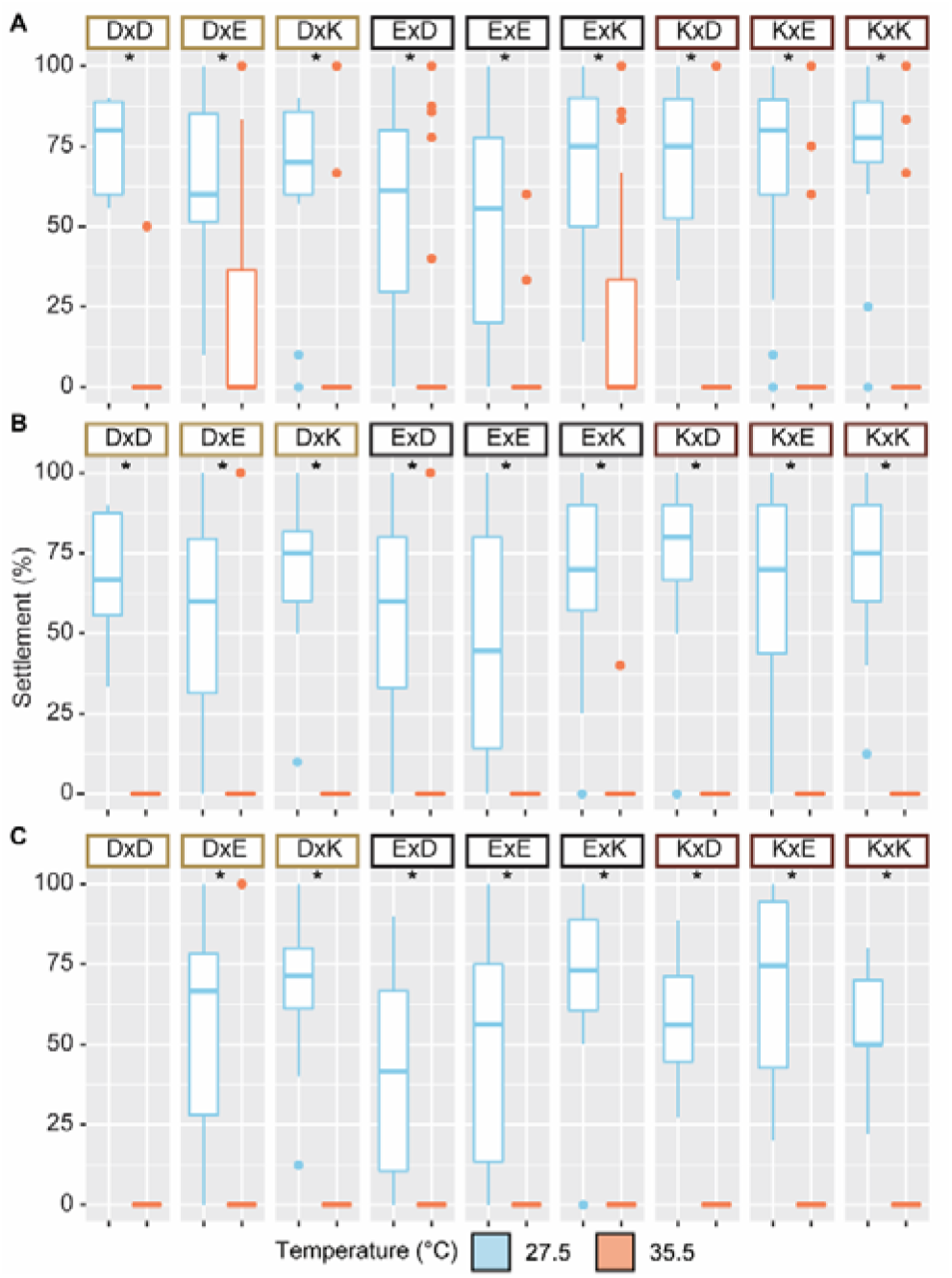
Larval settlement under experiential stress. Percentage of median settlement of n = 69 different *Acropora tenuis* lineages of larvae exposed to control (blue) or heat stress (red) conditions after 17 h (A), 24 h (B) and 48 h (C). The coloured boxes around the crosses indicate the source reef of the maternal colonies: Davies (tan), Esk (black), and Keppels (maroon). The Wilcoxon’s test was performed to analyze statistical differences

There was a significant effect of temperature treatment (F_1,1633_ = 887.50, *P =* 2.00×10^−16^) and heat exposure time (F_3,1631_ = 120.70, *P =* 2.00×10^−16^) on the percentage of settled larvae and a significant interaction between temperature treatment and heat exposure time (F_3,1564_ = 225.42, *P =* 2.00×10^−16^), temperature treatment and population level cross (F_8,1564_ = 5.30, *P =* 1.39×10^−6^). There was no significant interaction between heat exposure time and population-level cross (F_24,1564_ = 1.06, *P =* 3.86×10^−1^) or between temperature treatment, heat exposure time and population-level cross (F_23,1564_ = 1.26, *P =* 1.82×10^−1^). This significant decrease in settlement at 24 and 48 hours occurred regardless of larval cross (F_8,1626_ = 1.583, *P =* 1.25×10^−1^) or reef of origin of the maternal coral (F_2,1632_ = 0.133, *P =* 8.75×10^−1^).

After 17 hours of temperature incubation, a median value of 0.00 % larval settlement was observed for all crosses in the heat treatment and was significantly lower compared to the control temperature (median 70.00 %) across all larval lineages (Wilcoxon’s test, *P* < 0.001; **Fig. 3A**). In the control treatment, the median settlement percentage was highest in DaviesxDavies and KeppelsxEsk, with a median of 80.00 % settlement. The lowest median percentage settlement was observed in the EskxEsk crosses, with 55.50 % settlement and correspondingly high variability.

After 24 hours (**Fig. 3B**), median settlement remained at 70.00 % in the control treatment and ~0.00 % in the heat treatment, with all crosses settling significantly less at heat compared to controls (Wilcoxon’s test, *P* < 0.001). At this time point, KeppelsxDavies showed the highest percentage settlement (80.00 %) at control, followed by DaviesxKeppels and KeppelsxKeppels (75.00 %). Finally, after 48 hours (**Fig. 3C**), the median settlement percentage in control treatment dropped slightly to 66.67 %, where KeppelsxDavies crosses again exhibited the highest larval settlement at control (75.00 %). At heat, the only cross to settle included some of the larval replicates within DaviesxEsk. Finally, some crosses also experienced mortality of recent recruits from 17 to 48 hours (DaviesxDavies), whilst others continued to settle (DaviesxEsk).

### Adult responses to heat stress at the population level

#### Bleaching

There is a significant effect of temperature treatment (**Fig. 4A**; negative binomial generalized linear model-nbglm, *P* = 2.20×10^−16^) and population origin (nbglm, *P* = 3.56×10^−15^) on the median bleaching score of coral fragments after 16 days. At the control temperature, fragments sourced from Davies recorded a mean colour score of 4.75 ±0.14 (mean ±SE; median = 5.00) whilst fragments sourced from Keppels and Esk were 1.85 ±0.25 (median = 2.00) and 1.35 ±0.39 (median = 0.00), respectively (**Fig. 4A**). At heat, fragments from all populations bleached heavily (all mean scores <1.00, all median = 0.00). Relative to the control temperature, Davies fragments bleached the most (median percentage change 87.41 ±5.32 % decrease in bleaching category), followed by Keppels (62.96 ±37.04 %), and then Esk fragments (28.57 ±28.57 %). Pairwise comparisons showed that bleaching score was significantly different between the control and heat treatments in Davies (mean Tukey’s test, *P* < 0.001, median Wilcoxon *P* = 1.80×10^−15^), Esk (Tukey’s test, *P* < 0.001, median Wilcoxon *P* = 0.11), and Keppels (Tukey’s test, *P =* 0.15, median Wilcoxon *P* = 1.50×10^−2^).

**Figure 4.**
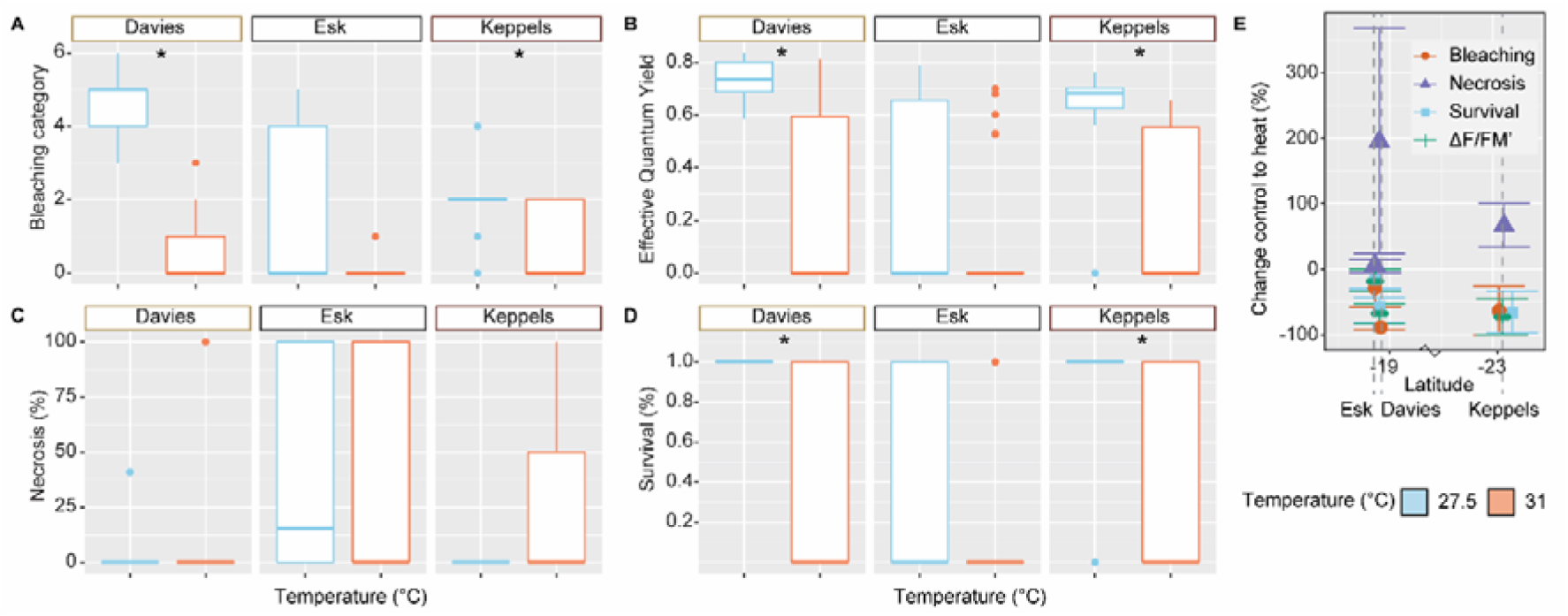
Median physiological responses in adult corals exposed to control and heat stress temperatures: Bleaching category core (A), Effective quantum yield (B), percent necrosis (C), and percent survival (D) of adult *Acropora tenuis* fragments. Colonies were collected from three sites on the Great Barrier Reef: Davies reef (tan outlines in A-D), Esk reef (black) and Keppels reef (maroon). Mean percent change in each response by latitude of collection site (E). The Wilcoxon’s test was performed to analyze statistical differences.

#### Photophysiology

The photophysiological responses, as measured by effective quantum yield (ΔF/FM’), showed that ΔF/FM’ significantly decreased in the heat compared to control treatment (**Fig. 4B**, lm, *P* = 1.09×10^−15^) and by population origin (lm, *P* = 6.80×10^−8^). Davies and Keppels fragments reported the highest yields at control temperatures (mean and median ΔF/FM’ >0.60), whereas Esk fragments were extremely low (0.12 ±0.05, median = 0.00). Mean ΔF/FM’ recorded from fragments in the heat treatment were highest in Keppels (0.22 ±0.07, median = 0.00), Davies (0.20 ±0.05, median = 0.00), and finally Esk (0.12 ±0.05, median = 0.00) but very variable overall. Relative to control temperatures, pairwise comparisons showed that only Keppels and Davies fragments significantly decreased ΔF/FM’ (Tukey’s test, *P* < 0.001 for both, median Wilcoxon = 2.40×10^−4^ for Keppels and *P* = 2.70×10^−12^ for Davies) but not Esk (Tukey’s test, *P =* 5.68×10^−1^, median Wilcoxon = 0.16).

#### Necrosis

Partial mortality was assessed as the percentage of necrotic tissue relative to each fragment (**Fig. 4C**). Population origin had a significant effect on percentage necrosis (lm, *P* = 3.25×10^−8^). There was no significant difference in percentage necrosis due to temperature treatment (lm, *P* = 5.50×10^−2^). At the control temperature, Davies and Keppels corals showed no to very little necrosis (median = 0.00 %), whereas Esk fragments were slightly necrotic (median 15.29 %). After 16 days at heat, fragments sourced from Esk lost on average, approximately 46.04 ±9.89 % (median = 0.00 %) of their tissue per fragment, compared to Keppels (26.67 ±11.82 %, median = 0.00 %) and Davies fragments (12.20 ±5.17 %, median = 0.00 %). Pairwise comparisons showed that at the control temperature, Esk lost, on average significantly more tissue compared to Davies (Tukey’s test, *P* < 0.001) and Keppels (Tukey’s test, *P* = 3.39×10^−3^). In the heat treatment, Esk also experienced significantly more necrosis compared to Davies (Tukey’s test, *P* = 2.76×10^−3^). All other population comparisons were not significantly different (Esk-Keppels, Tukey’s test, *P =* 5.35×10^−1^, Keppels-Davies, Tukey’s test, *P =* 7.43×10^−1^). Relative to control temperatures, the median percent necrosis was not significantly different for any population (median Wilcoxon *P* = 0.09, 0.91, 5.30×10^−2^ for Davies, Esk, Keppels).

#### Survival

Temperature showed significant effect on survival (binomial lm, *P =* 1.19×10^−8^). In the control treatment, survivorship was highest in the Keppels fragments (median = 1.00), followed by Davies (median = 1.00) and Esk (median = 0.00) (**Fig. 4D)**. In the heat treatment, survivorship was highest in the Keppels fragments (40.00 ±13.09 %, median = 0.00), followed by Davies (31.71 ±7.36%, median = 0.00) and Esk (20.00 ±8.16 %, median = 0.00). Compared between heat and control temperatures, fragments sourced from Davies and Keppels significantly decreased in survivorship (Tukey’s test, *P* < 0.001 and 4.98×10^−3^, median Wilcoxon = 5.10×10^−3^ and 1.40×10^−10^, respectively) but not Esk fragments, which also survived poorly at the control temperature treatment (Tukey’s test, *P =* 7.55×10^−1^, median Wilcoxon = 0.25). Given the significant population effect of temperature treatment for bleaching score, ΔF/FM’, and necrosis (but not survival), the relative differences in responses were also calculated and compared across the latitudinal gradient of adult origin (**Fig. 4D**). There was no trend in performance across traits by latitude.

### Predicting adult and offspring responses using gradients of selection

#### Adult responses

When survival was averaged at the population level, almost all adult corals collected from Davies exhibited approximately equivalent survival at the control treatment, but when exposed to the heat treatment, the population responses were generally bimodal, where some individuals within each population exhibited high survival and others low survival (**Fig. 5A**). Corals collected from Esk and the Keppels also exhibited this bimodal response for both the heat and control temperatures. For both Davies and Keppels corals, the mean survival at the control treatment was high and a roughly equal decrease in survival at heat. Although Esk corals had much lower survival at the control temperature, their overall mean survival at heat was roughly equivalent to Davies and Keppels corals, suggesting that the selection differential for survival at heat in each population was roughly equivalent (horizontal dashed black lines), defined as the difference between a selected phenotypic scope (triangle) and the mean percent survival at heat (vertical dashed red line).

**Figure 5.**
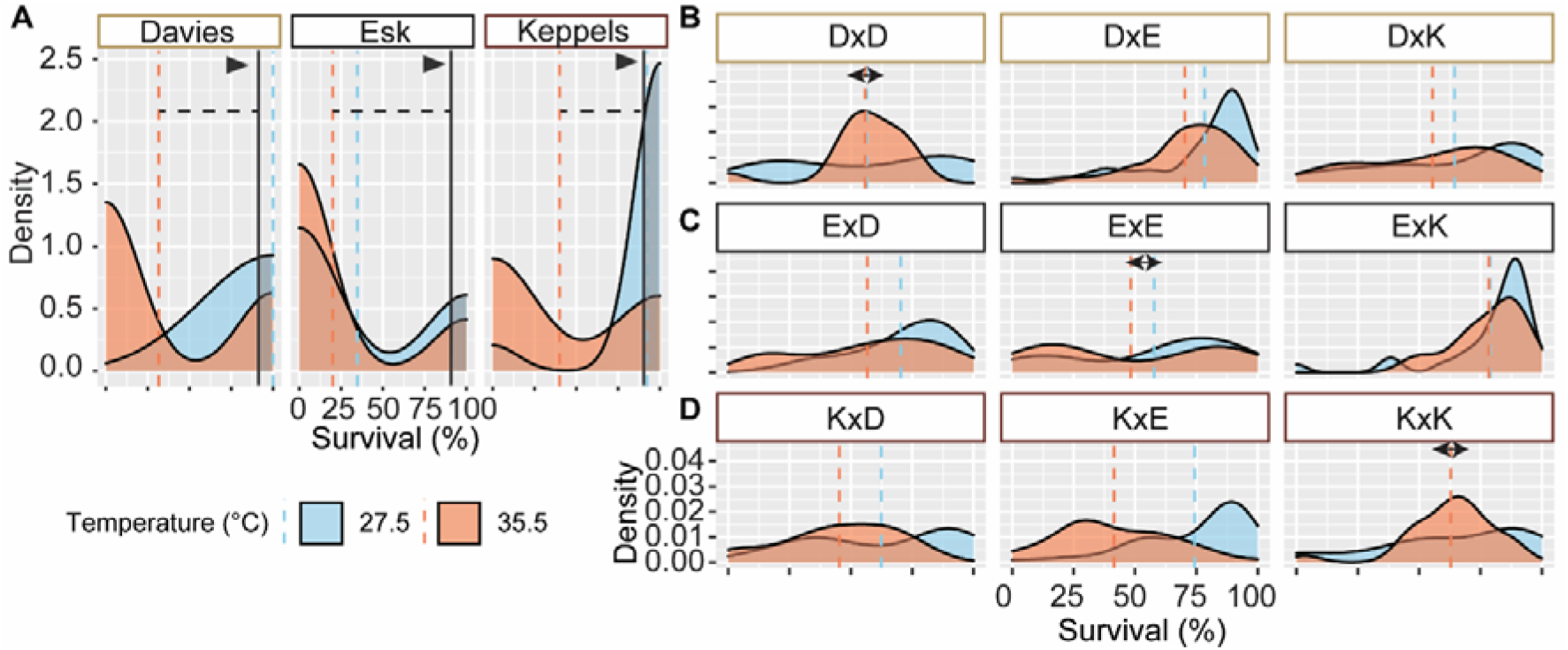
Percent survival density plot of the number of adult colonies (y) from each population (A) and offspring of each population cross (B-D). Arrows indicated the selected phenotypic scope. Horizontal dashed lines represent the selection differential for each population. Vertical dashed lines represent mean percent survival per population. The coloured boxes around the crosses indicate the source reef of the maternal colonies: Davies (tan), Esk (black), and Keppels (maroon). Median survivorship data is presented in Figures 2 and 4D, whereas survival data here is presented as the number of individual colonies per percent survival value. Survival at ambient and hot are shown together to aid in comparison.

#### Comparisons of purebred and hybrid larval responses to adults

Mean percent survival for larvae in the control treatments were higher (56.44 – 78.92 %) compared to at heat (41.48 – 78.40 %). As expected, based on similar adult selection differentials (**Fig. 5A**), purebred larvae from DaviesxDavies, EskxEsk, and KeppelsxKeppels responded similarly in the breadth of larval responses between control and heat (specifically, a small difference between treatment responses). Alternatively, interpopulation crosses differed in their responses, in which KeppelsxDavies and KeppelsxEsk had the largest difference in mean survival between control and heat (17.14 % and 32.84 %) and EskxKeppels the smallest (0.52 %).

The DaviesxDavies purebreds exhibited a 56.44 ±8.77 % mean survival at heat, almost 2x greater survival than the Davies adults at heat (31.71 ±7.36 %; **Fig. 5B**). DaviesxDavies larval crosses demonstrated high variability (e.g. flat distribution) responses in survival at control, compared to a unimodal response in heat, although the mean survival was roughly equal 56.44 ±8.77 % and 55.78 ±4.71 % respectively, dashed lines). When Davies eggs were crossed with sperm from the other central reef, heat tolerance increased by 14.46 % compared to purebred. DaviesxEsk larvae exhibited unimodal responses for both temperatures with average survival at 78.38 % at control and 70.24 % at heat. Heat tolerance was slightly less for DaviesxKeppels larvae (55.41 %) compared to DaviesxDavies, where larval responses were more variable but still unimodal.

The EskxEsk larval purebreds exhibited 48.29 ±7.15 % survival at heat, ~2.5x greater survival than the Esk adults at heat (20.00 ±8.16 %; **Fig. 5C**). EskxEsk larvae at control and heat treatments were relatively flat, demonstrating that both treatments consisted of crosses with high and low survival. When Esk eggs were crossed with sperm from the other reefs, heat tolerance increased by 8.40 % and 30.11 % in EskxDavies and EskxKeppels, respectively, relative to EskxEsk. EskxDavis larval survival responses were weakly unimodal with mean survival at 70.29 ±3.47 % and 56.68 ±4.61 % for control and heat. Alternatively, EskxKeppels larvae exhibited strongly unimodal survival responses at 78.92 ±3.82 % at control and 78.40 ±2.59 % at heat.

The KeppelsxKeppels purebreds exhibited 62.84 ±3.73 % survival at heat, ~1.5x greater survival than the Keppels adults at heat (40.00 ±13.09 %; **Fig. 5D**). KeppelsxKeppels larval responses formed relatively flat distributions in the control treatment to unimodal responses in the heat. When Keppels eggs were crossed with sperm from the other reefs, heat tolerance decreased by 17.59 % and 21.35 % in KeppelsxDavies and KeppelsxEsk, respectively. In KeppelsxDavies, the distribution was flat wide, indicating variation in survival. The KeppelsxEsk density plot for larvae at the control treatment exhibited a unimodal peak in survival, as well as in the heat treatment.

In the heat stress treatment when larvae were grouped by the population identity of the maternal colony, larvae produced from Davies and Keppels had selective landscapes that were wider compared to Esk larvae (**Fig. 6A-C**), although Davies and Esk had overall higher selection coefficients compared to Keppels larvae (S= Esk: 60 –70, Davies: 54 to 70, Keppels 50 – 60; **Fig. 6A-C**). Combined with estimated narrow-sense heritability estimates, these differences resulted in overall higher selective responses (R) for Davies and Esk (i.e., more values between R = 40 – 60) compared to Keppels offspring (**Fig. 6D-F**).

**Figure 6.**
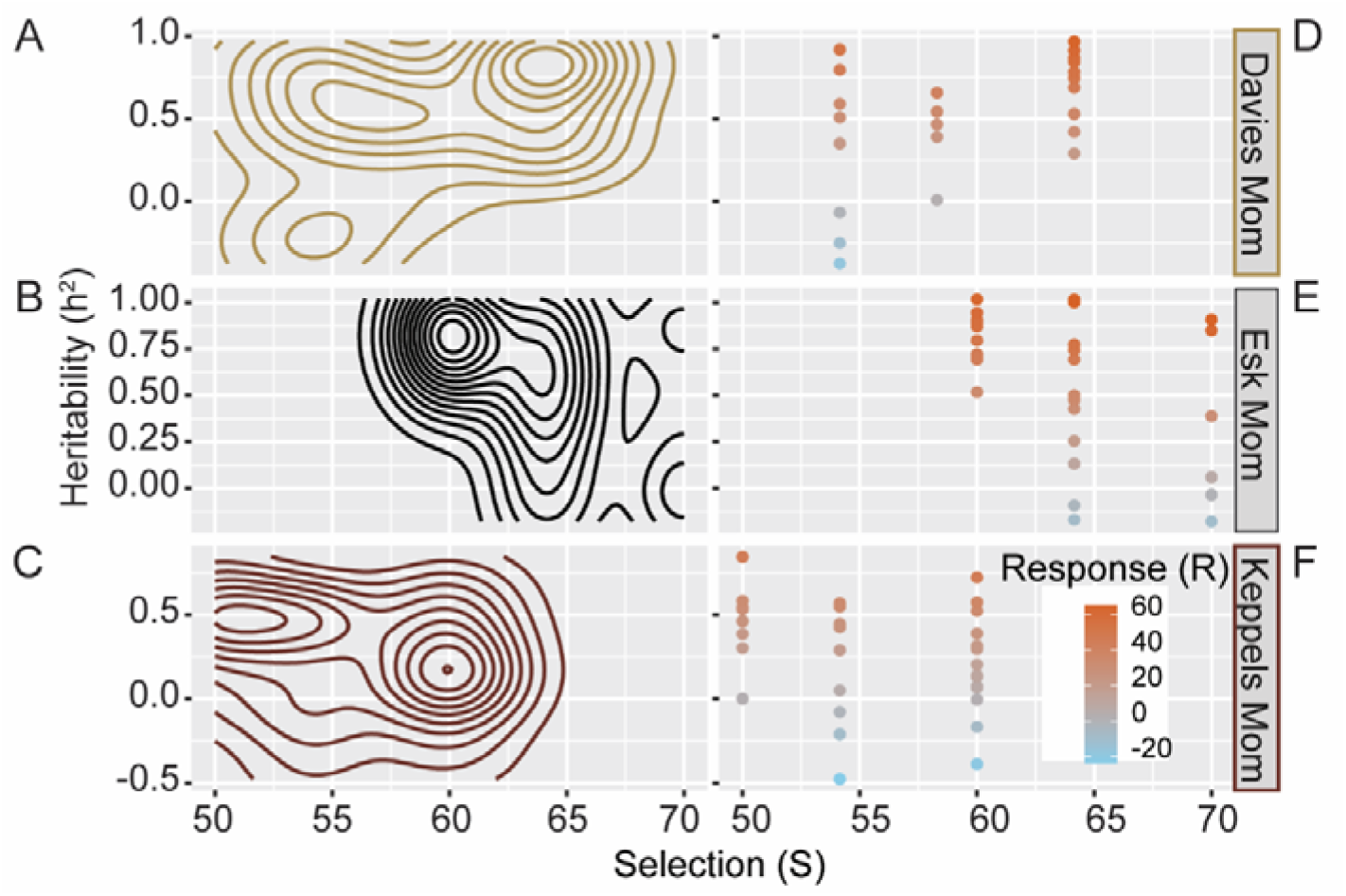
Adaptive landscape of selection for heat tolerance. Modelled selective values (S) and narrow-sense heritability (h^2^) values of larval survival at heat grouped by the population identity of the maternal colony (A-C). Estimated response to selection (R) for h^2^ and S (D-F).

## DISCUSSION

Mean maximum sea surface temperatures are expected to increase by between 2 – 4 °C by 2100 globally (IPCC, 2014). Without adaptation, this will likely exceed the thermal thresholds of corals. Understanding the underlying adaptive capacity of wild populations is therefore critical to forecasting species persistence. Moreover, various conservation strategies are being considered worldwide to help corals withstand increasing ocean temperatures whilst carbon emissions are curbed (National Academies of Sciences, Engineering, 2019). This includes the introduction of more heat-tolerant offspring produced from selective breeding methods onto cooler reefs to prepare them for warming (Quigley et al., 2018), a method that will also increase the genetic diversity on reefs, fuel for which natural selection can act. Quantifying the feasibility for enhancing corals’ ability to survive further ocean warming is therefore vital for the conservation of the world’s coral reefs.

### Little variation in adult physiological responses to heat stress across three GBR populations

Phenotypic variation in organisms drives the capacity for plastic, adaptive responses to environmental pressure. This variation may be underpinned by genetic variation or by responses mediated by non-genetic mechanisms, like changes in the microbiome (e.g. bacteria or Symbiodiniaceae), in which algal symbiont assemblages may shape corals’ responses to heat stress (Berkelmans and van Oppen, 2006). Understanding the scope for phenotypic variation to heat stress at the adult stage is essential to evaluating the scope for heritable diversity of heat tolerance at later life stages in corals.

Overall, there was no difference in heat tolerance of adult southern Keppels corals compared to central Davies corals, with both populations suffering similar percent necrosis, drops in photophysiology, and lowered survival under heat stress. However, the relative change in bleaching was greater for Keppels corals. The magnitude of bleaching was also more severe in Davies compared to Keppels corals. *Acropora tenuis* in both central and southern reefs generally hosts dominant abundances of *Cladocopium* (Rocker et al., 2017; Ulstrup and van Oppen, 2003), which could contribute to the similarity in their physiological performance, whereas, symbionts from Davies reef or the host corals themselves may have lower initial tolerances but are able to recover and survive equally well. Taken together, these results suggest that the absolute heat tolerance of both populations was roughly equal. Finally, adult coral fragments from the Esk population in heat treatment exhibited the highest bleaching severity, lowest effective quantum yield, highest percentage necrosis, and lowest survivorship. It should be noted that although fragments sourced from the Esk population lost the greatest overall percentage of tissue (necrosis) per population, fragments in the control treatment were also highly necrotic, suggesting a compromised health state of corals from this population, also reflected in the low survival and photosynthesis in controls. Both Davies and Esk corals were collected during the sample trip with the same level of handling. This suggests that transport issues were not the cause of their diminished health state but instead point towards population-level differences between these corals.

The high overall fitness of adult Keppels corals under heat stress was surprising. This population reported the lowest bleaching severity, highest effective quantum yield, and highest survival at heat. Although enhanced heat tolerance in corals is generally attributed to corals from warmer reefs exhibiting higher upper thermal thresholds (Berkelmans, 2002; Howells et al., 2012; Ulstrup et al., 2006), the enhanced performance of Keppels corals may be attributed to the greater variability in their local thermal regime. The increase in tolerance may also partly be attributed to the coral symbionts (Howells et al., 2012; Thomas et al., 2018; Ulstrup et al., 2006), and their interaction with host genetics (Dixon et al., 2015; Smith-Keune and van Oppen, 2006; Thomas et al., 2018), in which complex holobiont interactions influence the overall heat stress-responses via gene regulation, symbiont density control and assemblage shuffling (Cunning and Baker, 2020; Yuyama et al., 2018). Our results indicate that the control of heat tolerance is complex and that many factors, including local thermal regime, likely play a role.

### Minimal improvement in larval settlement due to selection for heat tolerance

Previous breeding experiments have demonstrated the transfer of increased offspring survival from parents sourced from warm reefs when reproductively crossed with cooler reefs (Dixon et al., 2015), or at least one parent from warmer reefs (Quigley et al., 2020b), suggesting genetic contribution to offspring. In this study, larval survival was high in the crosses whose mother sourced from either Esk or Davies. However, it is currently unknown whether an increased propensity for settlement at high temperatures is also transferable using colonies sourced from warmer reefs to achieve an enhancement in settlement success. Although settlement is a heritable trait under control conditions (h^2^ = 0.49; Meyer et al., 2009), the overall heritability is low relative to other fitness-related traits. Moreover, it is well known that settlement in corals is negatively impacted by heat. For example, early life-stage *Acropora tenuis* settlement decreased by 100 % when exposed to +5 °C over ambient temperature (Humanes et al., 2016), and by 55 % when combined with suspended sediments treatment (Humanes et al., 2017). *Diploria strigose* larvae demonstrated a decrease in the settlement behaviour at temperatures exceeding 30 °C compared to control (Bassim and Sammarco, 2003) and *Acropora palmata* settlement decreased by 25 % at 31.5 °C compared to 28 °C (Randall and Szmant, 2009). The lack of strong differences in settlement success between the crosses here may be reflected in the roughly equal heat tolerance of both Davies and Keppels corals, suggesting that both populations are roughly equivalent in tolerance and therefore did not produce strong differences in settlement of larvae. Combined, these previous results suggest that selection should act on this important trait over time if oceans continue to warm.

The results from this study show that, when exposed to heat, larvae from all crosses significantly decreased in their settlement behaviour relative to the control temperature. During periods of increased temperature, the cellular processes within larvae become compromised, including disruption in the repair of cellular proteins and enzymes (Negri et al., 2007). Consequently, abnormalities develop, and the rate of cell cleavage rapidly increases. This cellular impairment could prevent the transition from larvae to recruit by preventing attachment to substrata or an increase in hypersensitivity during transition phases of larvae to polyp (Randall and Szmant, 2009). The reduction in larval settlement could also be a consequence of energy deficiency. As cellular proteins unfold and aggregate, HSP70, a heat shock protein that refolds degrading proteins (Daugaard et al., 2007) has been found to be upregulated in *Acropora millepora* larvae to maintain normal cell function (Rodriguez-Lanetty et al., 2009). This requires vast amounts of energy, that could potentially otherwise be used for settlement. Higher respiration rates at heat also increase metabolic activity (Edmunds et al., 2001) which, in turn, increases the amount of food required to maintain these elevated levels. These factors could all contribute to the reduced settlement of larvae measured here.

Overall, although our results showed a significant decrease in larval settlement at heat compared to control temperatures, this did not correspond to reef of origin. Our findings only weakly allude to the potential for selectively bred coral larvae to settle at higher temperatures. For example, while our results do not demonstrate a significant increase, there was a higher percentage of settled larvae whose mother coral was from either Esk or Davies (the central, warmer, inshore reefs in this study) compared to those with a mother coral from the southern, cooler Keppels reef, in contrary of adult’s response in which Keppels had higher overall fitness than Davies and Esk. Previous research suggests mitochondrial DNA (mtDNA) plays a large role in the thermal resistance of corals, alluding to a high maternal effect on the heat tolerance of coral offspring (Dixon et al., 2015; Quigley et al. 2020). There was also some variation in settlement within population crosses under heat, demonstrating the potential for plasticity. This aligns with information that corals found in warmer environments or with high daily temperature variability have greater genetic plasticity (Kenkel and Matz, 2017) which can be passed onto offspring. However, the findings presented here are preliminary and it appears that the maximum thermal limits of parental adult corals and the resulting larvae are not indicative of settlement success. As warming increases in severity, these processes may be the first to be disrupted (Radchuk et al., 2019), and assessing the impacts on these and other fundamental processes like recruitment will become increasingly important.

The lack of a significant effect of parental colony from warmer reefs to enhance settlement at high temperatures may be due to either experimental or biological factors. The high temperatures chosen here may have surpassed the corals’ settlement behaviour, in which temperatures exceeding 35.5 °C were extreme compared to the mean monthly maximums of these sites (all 24 - 27 °C), resembling short-term acute heat stress temperature range (Grottoli et al., 2021; McLachlan et al., 2020). However, the experimental temperature of many studies does not exceed +5 °C above the control temperature (Humanes et al., 2016; Humanes et al., 2017; McLachlan et al., 2020; Quigley et al., 2020b). Hence, our result could reflect the contribution of higher-than-threshold temperature treatment. Alternatively, environmental factors may contribute a greater influence in determining settlement compared to host genetics. Specifically, settlement deficiency at high temperature may be driven by the disruption of the microbial biofilm needed to induce metamorphosis and settlement, where it is well established that settlement is induced by the presence of CCA and microbial biofilms (Webster et al., 2004). During periods of increased temperature, chemical cues released by CCA can be weakened and microbial cells present in biofilms (on the CCA) can become damaged (Randall and Szmant, 2009). Therefore, this reduction in biochemical cues could be the main contributing factor to the reduction in larval settlement at higher temperatures.

In summary, larvae from intra- and interpopulation crosses demonstrated a general inability to settle at high temperatures, albeit extreme, suggesting that improvement in larval settlement responses due to selection for heat tolerance may be challenging due to the competing influence of environmental effects. Combined with information that behavioural or morphological traits generally respond less to selection compared to life-history traits (Mousseau and Roff, 1987) and the potential governing importance of environmental factors (e.g. bacterial communities), this suggests that the enhancement of this trait under heat stress may require the selection through microbial community contribution more than through processes targeting host genetics.

### Predicting responses to heat using selection differentials and gradients of selection

Genetic variation underpins the potential and speed for adaptation through natural selection (Falconer and Makcay, 1996). Warming influences traits differentially, with morphological traits generally less impacted compared to phenological traits (Radchuk et al., 2019). Survival was chosen as the trait of interest to examine gradients of selection given the importance of survival and other life-history traits compared to behavioural or morphological traits (Mousseau and Roff, 1987). Measures such as narrow-sense heritability (h^2^), selection (S), and responses to selection (R) are useful for quantitively assessing organisms’ ability to respond to their environment, especially future stressors. Narrow-sense heritability ranges from 0 – 1, where 0 is indicative of no genetic contribution to trait variance and 1 is complete dominance of genetics in determining trait variance (Falconer and Makcay, 1996). Measurements derived from corals suggest that they do have a strong underlying capacity to respond adaptively to heat either through host genetics (h^2^ = mean: 0.86, range: 0.48 - 0.93) (Dixon et al., 2015; Dziedzic et al., 2019; Kirk et al., 2018; Quigley et al., 2020b) or through changes to their symbiont communities (Quigley et al., 2018). It might be expected that heat stress would elicit directional selection through differential mortality of adults, resulting in the survival of a subset of phenotypes at one end of the phenotypic distribution. However, the bimodal responses in survival curves of coral adults suggest that heat stress manifests as disruptive selection, which may explain the high variability of heat responses across the numerous offspring lineages. Although the underlying mechanisms are unknown here, the drivers of bimodality may be linked to biochemical complexity (Rezende and Bozinovic, 2019).

Selection differentials depend on the heritability of the trait, where heritability is generally equal to the slope of the Response over Selection, as determined by the breeder’s equation. Interestingly, the selection differentials at heat (i.e. the intensity of adaptive responses) were similar across the three populations of adult corals. This mirrors the similar physiological responses of the adult corals to heat stress. The selection differential between survival at high temperatures can be described as the difference between the mean value measured (Davies and Keppels = ~40 - 30 %, and Esk ~25 %) and the desired mean value (e.g. ~90 %). Hence, the selection differential was about 55 % (90 minus 35 %) across these three populations, which represents a desired 63 % increase in potential trait enhancement. When translated to estimated adaptive larval responses, these varied by population cross. In comparison, responses to selection in well-studied systems like aquaculture species, including fish and shellfish, averaged about ~13 % for growth and 4.9 % for survival (Gjedrem and Rye, 2018). Furthermore, our comparative analysis between adult and offspring responses suggests a divergence between adult population mean responses and interpopulation offspring crosses. The divergence between thermal responses of different life-stages within species has been demonstrated in other marine organisms like mollusks (Truebano et al., 2018), in which larval forms are often more or less vulnerable to heat compared to adults. In this case, mollusks reported a 1.7 - 2.1x difference in responses between larvae and adults, which mirrors reports seen in other invertebrates like brine shrimp (2.7 - 4.9x; Norouzitallab et al., 2014), copepod (Tangwancharoen, 2014), and other invertebrates (Pandori and Sorte, 2019). Finally, adaptive response surfaces revealed that when larvae were grouped by their maternal populations, Davies and Esk offspring generally had higher selective responses (R) compared to Keppels offspring. Taken together, this suggests that although adult populations may respond similarly to heat, the overall potential of offspring responses to selection in warmer populations of corals from Davies and Esk is greater compared to cooler populations in the Keppels. These findings have important implications for forecasting the impacts of climate change on wild populations of corals and for the development of novel conservation tools like the assisted evolution of at-risk populations.

## Conclusions

As climate change accelerates ecosystem change, critical information on how important fitness traits will vary in the future is essential to move from understanding impacts to predicting and forecasting those impacts. This comparative physiological dataset across different life-history stages in one important coral species provides key mechanistic and adaptive insights into how corals may function under heat stress caused by warming oceans, and in particular, the heritability of heat tolerance.

## Supporting information

R scripts using in the study

Data sets using in the study

Source file of figures in the EPS format

Responses to comments from journal's reviewer

## Acknowledgements

The authors would like to thank the Traditional Owners from whose Sea Country these corals were collected. Specifically, we would like to thank the Woppaburra, Manbarra, Bindal, and Wulgurukaba Traditional Owners. Corals were collected under the following permit numbers: G12/35236.1 (Davies and Esk) and G19/43024.1 (Keppels) to the Australian Institute of Marine Science. We would like to thank Andrea Severati, Line Bay, Christine Giuliano, and the spawning collections field crews in the central and southern Great Barrier Reef for assisting in the coral colony collections.

## Footnotes

### Author contributions

Conceptualization and Methodology: K.M.Q.; Formal analysis: P.W.; K.M.Q.; Investigation: P.W., R.C., A.M., H.K., J.R., K.M.Q; Resources: K.M.Q.; Data curation: P.W., K.M.Q. Writing – original draft: K.M.Q., R. C., J.R.; Writing - review & editing: P.W., R.C., A.M., K.M.Q.; Supervision: K.M.Q.; Funding acquisition: K.M.Q.

### Data availability

Physiological data and analysis code is available at https://github.com/LaserKate/KeppelsAGF19

