## Supplementary material for "Predicting selection-response gradients of heat tolerance in a widespread reef-building coral": R scripts using in the study: Adult_density_plot.R

```
####################################
#import dataset
AGF19_Adult<-read.csv("D:/PhD/Journal/adult/Adult_AGF19_Tfinal_response.csv")
library(dplyr)
library(lattice)
library(plyr)
library(Rmisc)

####################################
#plot density
library(ggplot2)
library(ggpubr)
adult_surv<-ggplot(AGF19_Adult, aes(x=Survival, fill=Temperature)) +
  geom_density(alpha=0.6)+
  scale_fill_manual(name="Temperature" ,values = c("31C"="coral","27.5C"="skyblue"))+
  facet_wrap(.~ Population, ncol=3)+
  ggtitle("Adult Survival")+
  theme(plot.title = element_text(size = 14, family = "Tahoma", face = "bold"),
        text = element_text(size = 12, family = "Tahoma"),
        axis.title = element_text(face="bold"),
        axis.text.x=element_text(size = 11))

#adult_survival+geom_vline(aes(xintercept = grp.mean, color = Population), data = AGF19_Adult, linetype = "dashed")

####################################
#compute mean adult survival
adult_surv_mean<-summarySE(AGF19_Adult, measurevar="Survival", groupvars=c("Population", "Temperature"), na.rm=TRUE)
adult_surv_mean
adult_surv_mean %>% arrange(Temperature,Population)
write.csv(adult_surv_mean,"D:/PhD/Journal/adult\\adult_surv_mean.csv", row.names = FALSE)

#add mean line
adult_surv+
  geom_vline(aes(xintercept = Survival, color = Temperature), data = adult_surv_mean, linetype = "dashed")+
  scale_color_manual(values = c("31C"="coral","27.5C"="skyblue"))
#geom_text(data = adult_surv_mean, aes(label = round(Survival, digits=2), y = Survival + 2, x = Survival - 0.02))
```
