## Supplementary material for "Predicting selection-response gradients of heat tolerance in a widespread reef-building coral": R scripts using in the study: Adult_latitude_change_plot.R

```
#plot changes per latitude
AGF19_Adult<-read.csv("D:/PhD/Journal/adult/AGF19_Tfinaldataa.csv")
#
bl<-summarySE(data=AGF19_Adult, measurevar="Bleaching", groupvars=c("Genotype", "Temperature", na.rm=TRUE))
yii<-summarySE(data=AGF19_Adult, measurevar="YII", groupvars=c("Genotype", "Temperature", na.rm=TRUE))
nec<-summarySE(data=AGF19_Adult, measurevar="Perc.necrosis", groupvars=c("Genotype", "Temperature", na.rm=TRUE))
surv<-summarySE(data=AGF19_Adult, measurevar="Survival", groupvars=c("Genotype", "Temperature", na.rm=TRUE))
#
df <- read.csv("D:/PhD/Journal/adult/latitude_change.csv")
dat<-summarySE(data=df, measurevar="Percentage", groupvars=c("Traits", "Latitude", "Population", na.rm=TRUE))
write.csv(dat,"D:/PhD/Journal/adult\\Adult_latitude_change_result.csv", row.names = FALSE)
#
library(Rmisc)
library(ggplot2)
library(scales)
#
p <- ggplot(dat, aes(x=Latitude, y=Percentage, group=Traits, color = Traits))+
  geom_point(aes(shape=Traits), size = 2, na.rm=TRUE)+
  geom_errorbar(aes(ymin=Percentage-se, ymax=Percentage+se, color = Traits), width = 0.5)+
  scale_color_manual(values = c("Bleaching"="#D76127","YII"="#199E77","Necrosis"="#7570B3", "Survival"="#87CCE9"))+
  scale_x_reverse()+
  ylab("Percentage of changes from ambien to heat treatment")+
  ggtitle("Changes between two treatments")+
  xlab("Latitude")+
  theme_gray() +
  theme(plot.title = element_text(size = 14, family = "Tahoma", face = "bold"),
        text = element_text(size = 12, family = "Tahoma"),
        axis.title = element_text(face="bold"),
        axis.text.x=element_text(size = 11, angle = -45),
        axis.title.y = element_text(size = 8),
        panel.border= element_rect(color="black", fill = NA))

p
```
