## Supplementary material for "Predicting selection-response gradients of heat tolerance in a widespread reef-building coral": R scripts using in the study: Adult_median_plot.R

```
#
library(dplyr)
library(plyr)
#
AGF19_Adult<-read.csv("D:/PhD/Journal/adult/Adult_AGF19_Tfinal_response.csv")
write.csv(p_meds,"D:/PhD/Journal/adult\\median_necrosis.csv", row.names = FALSE)
#
library(ggplot2)
library(ggpubr)
#
b_meds <- ddply(AGF19_Adult, .(Population, Temperature), summarise, med = median(Bleaching))

bleaching<-ggplot(AGF19_Adult, aes(x = Temperature , y = Bleaching)) +
  geom_boxplot(aes(color=Temperature))+
  facet_grid(.~Population)+
  scale_color_manual(values = c("31C"="coral","27.5C"="skyblue")) +
  ggtitle("A")+
  theme(plot.title = element_text(size = 14, family = "Tahoma", face = "bold"),
        text = element_text(size = 10, family = "Tahoma"),
        axis.title = element_text(face="bold", size = 12),
        axis.text.x=element_text(size = 10))+
  geom_text(data = b_meds, aes(y = med, label = med), 
            size = 2, vjust = -1.5)+
  stat_compare_means(aes(label = ..p.format..), size = 3, hjust = -1)
bleaching
#
y_meds <- ddply(AGF19_Adult, .(Population, Temperature), summarise, med = median(YII))

yii<-ggplot(AGF19_Adult, aes(x = Temperature , y = YII)) +
  geom_boxplot(aes(color=Temperature))+
  facet_grid(.~Population)+
  scale_color_manual(values = c("31C"="coral","27.5C"="skyblue")) +
  ggtitle("B")+
  ylab("Effective Quantum Yield")+
  theme(plot.title = element_text(size = 14, family = "Tahoma", face = "bold"),
        text = element_text(size = 10, family = "Tahoma"),
        axis.title = element_text(face="bold", size = 12),
        axis.text.x=element_text(size = 10))+
  geom_text(data = y_meds, aes(y = med, label = med), 
            size = 2, vjust = -1.5)+
  stat_compare_means(aes(label = ..p.format..), size = 3, hjust = -1)
yii
#
p_meds <- ddply(AGF19_Adult, .(Population, Temperature), summarise, med = median(Perc.necrosis))

nec<-ggplot(AGF19_Adult, aes(x = Temperature , y = Perc.necrosis)) +
  geom_boxplot(aes(color=Temperature))+
  facet_grid(.~Population)+
  scale_color_manual(values = c("31C"="coral","27.5C"="skyblue")) +
  ggtitle("C")+
  ylab("Percentage of necrosis")+
  theme(plot.title = element_text(size = 14, family = "Tahoma", face = "bold"),
        text = element_text(size = 10, family = "Tahoma"),
        axis.title = element_text(face="bold", size = 12),
        axis.text.x=element_text(size = 10))+
  geom_text(data = p_meds, aes(y = med, label = med), 
            size = 2, vjust = -1.5)+
  stat_compare_means(aes(label = ..p.format..), size = 3, hjust = -1)
nec
#
s_meds <- ddply(AGF19_Adult, .(Population, Temperature), summarise, med = median(Survival))

surv<-ggplot(AGF19_Adult, aes(x = Temperature , y = Survival)) + 
  scale_y_continuous(limits=c(0,1.1), breaks=c(0.2, 0.4, 0.6, 0.8, 1.0))+
  geom_boxplot(aes(color=Temperature))+
  facet_grid(.~Population)+
  scale_color_manual(values = c("31C"="coral","27.5C"="skyblue")) +
  ggtitle("D")+
  theme(plot.title = element_text(size = 14, family = "Tahoma", face = "bold"),
        text = element_text(size = 10, family = "Tahoma"),
        axis.title = element_text(face="bold", size = 12),
        axis.text.x=element_text(size = 10))+
  geom_text(data = s_meds, aes(y = med, label = med), 
            size = 2, vjust = -1.5)+
  stat_compare_means(aes(label = ..p.format..), size = 3, hjust = -1)
surv
#
ggarrange(bleaching, yii, nec, surv,
          ncol = 2, nrow = 2)
```
