## Supplementary material for "Predicting selection-response gradients of heat tolerance in a widespread reef-building coral": R scripts using in the study: Adult_response_plot.R

```
##Load your packages----
library(ggplot2)
library(nlme)
library(lme4)
library(plyr)

##read yourdata
AGF19_Adult<-read.csv("D:/PhD/Journal/adult/Adult_AGF19_Tfinal_response.csv")
head(AGF19_Adult)

#############################################################
#Run model with interaction of fixed factors
library(multcomp)
library(mvtnorm)
library(survival)
library(TH.data)
library(MASS)
AGF19_Adult$INT<-interaction(AGF19_Adult$Temperature, AGF19_Adult$Population)
#Now the object AGF19_Adult$INT within your dataset AGF19_Adult has a unique level for every combination of site and treatment. Write a new model for the purpose of post-hoc comparisons

#############################################################
##Bleaching is not normal data
#change D scale to numeric scale so you can take averages
## Bleaching D0 = dead
#Test random effects of tank and stick
Bleaching_model_Random<-glm.nb(Bleaching~Tank+Stick, data=AGF19_Adult, na.action=na.omit)
anova(Bleaching_model_Random)

#see if the data can be used with linear model
Bleaching_model1<-lm(Bleaching~Population+Temperature, data=AGF19_Adult, na.action=na.omit) 
summary(Bleaching_model1)
anova(Bleaching_model1)
plot(Bleaching_model1, 3)
qqnorm(Bleaching_model1)
#not suitable for lm

#generalized non binomial linear model
Bleaching_model1<-glm.nb(Bleaching~Population+Temperature, data=AGF19_Adult, na.action=na.omit)
anova(Bleaching_model1)
Bleaching_model1_glht<-glm.nb(Bleaching~-1+INT, data=AGF19_Adult, na.action=na.omit)
#Then, analyze post hoc comparisons
summary(glht(Bleaching_model1_glht, linfct=mcp(INT="Tukey")))

########################################### 
#YII
#Test random effects
YII_model_Random<-lm(YII~Tank+Stick, data=AGF19_Adult, na.action=na.omit)
anova(YII_model_Random)
plot(YII_model_Random, 1)
#no random effect

#linear model
YII_model1<-lm(YII~Population+Temperature, data=AGF19_Adult, na.action=na.omit)
anova(YII_model1)
plot(YII_model1, 2)
YII_model1_glht<-lm(YII~-1+INT, data=AGF19_Adult, na.action=na.omit)
#Then, analyze post hoc comparisons
summary(glht(YII_model1_glht, linfct=mcp(INT="Tukey")))

#############################################
#Necrosis
#Test random effects
Nec_model_Random<-lm(Perc.necrosis~Tank+Stick, data=AGF19_Adult, na.action=na.omit)
summary(Nec_model_Random)
anova(Nec_model_Random)
plot(Nec_model_Random, 2)
#no random effect

#linear model
Perc.nec_model1<-lm(Perc.necrosis~Population+Temperature, data=AGF19_Adult, na.action=na.omit)
anova(Perc.nec_model1)
plot(Perc.nec_model1, 2)
Perc.nec_model1_glht<-lm(Perc.necrosis~-1+INT, data=AGF19_Adult, na.action=na.omit)
#Then, analyze post hoc comparisons
summary(glht(Perc.nec_model1_glht, linfct=mcp(INT="Tukey")))

##############################################
#Survival
#Test random effects
Survival_model_Random<-glm(Survival~Tank+Stick, data=AGF19_Adult, family="binomial")
summary.glm(Survival_model_Random)
#no random effect

#generlizedd linear model
survival_model1 <- glm(Survival~Population+Temperature, data=AGF19_Adult, family="binomial") 
summary.glm(survival_model1)
survival_model1_glht<-glm(Survival~-1+INT, data=AGF19_Adult, na.action=na.omit) 
summary(glht(survival_model1_glht, linfct=mcp(INT="Tukey")))
```
