## Supplementary material for "Predicting selection-response gradients of heat tolerance in a widespread reef-building coral": R scripts using in the study: AGF19_temperature_plot.R

```
###################################Read in data
AGF19_Temps<-read.csv("D:/Master_IMBRSea/thesis at AIMS/Sat_eReefs_AGF2019Adult_temps.csv")
library(dplyr)
library(lattice)
library(plyr)
library(Rmisc)

###################################
#2019 Temperature
AGF19_SE_Temps<-summarySE(data=AGF19_Temps, measurevar="mean", groupvars=c("Month","Year", "Site.Name", na.rm=TRUE))

#
library(ggplot2)
library(ggpubr)

Temps_plot<-ggplot(AGF19_SE_Temps, aes(x = Month , y = mean, group = Site.Name)) +
  geom_line(aes(linetype=Site.Name, color=Site.Name))+
  geom_point(aes(color=Site.Name))+scale_x_discrete(limits = c(1,2,3,4,5,6,7,8,9,10,11,12))+
  geom_errorbar(aes(ymin=mean-sd, ymax=mean+sd, color=Site.Name), width=0.2)+
  ggtitle("AGF 2019 Temperature") +
  theme_bw() +
  theme(plot.title = element_text(size = 14, family = "Tahoma", face = "bold"),
        text = element_text(size = 12, family = "Tahoma"),
        axis.title = element_text(face="bold"),
        axis.text.x=element_text(size = 11))
Temps_plot
Temps_plot+scale_colour_manual(values = c("Keppels"="#6B322B","Davies"="#B0965A","Esk"="#020202"))

###################################
```
