## Supplementary material for "Predicting selection-response gradients of heat tolerance in a widespread reef-building coral": R scripts using in the study: Larvae_cumulative_hours.R

```
#import dataset
larv2019<-read.csv("D:/PhD/Journal/larvae/2019_Larvae_data_curated_JC.csv")
#
test <- larv2019 %>% dplyr::select(Timepoint_Master, Cumulative.hrs, Cone, Boat.ID,
                                   
                                   Wells, No.LarvaeInObserved, Percent_Survival_atend)

# test2 <- larv2019 %>% dplyr::select(Timepoint, Cumulative.hrs, Cone, Boat.ID, Wells,

#                                    No.LarvaeInObserved, Percent_Survival_atend) %>%

#   filter(Timepoint == "1")

test$unique.id <- paste(test$Cone, test$Boat.ID, test$Wells, sep='')

test$Cumulative.hrs <- as.numeric(test$Cumulative.hrs)

test <- test[order(test$unique.id, test$Cumulative.hrs),]


# plot % survival over time for each rep for dirty visual

check19 <- ggplot(data=test, aes(x=Cumulative.hrs, y=Percent_Survival_atend,
                                 
                                 group=unique.id)) +
  
  geom_line(size=0.001, alpha=0.4)
```
