## Supplementary material for "Predicting selection-response gradients of heat tolerance in a widespread reef-building coral": R scripts using in the study: Larvae_settlement_factor_script.R

```
AGF19_Larvae <- read.csv("D:/PhD/Journal/larvae/Larvae_AGF19_settlement.csv")
################################################
#Testing factors on settlement
mod1<-aov(SettledPercentage~TimePoint, data = AGF19_Larvae)
summary(mod1)

mod2<-aov(SettledPercentage~Treatment, data = AGF19_Larvae)
summary(mod2)

mod3<-aov(SettledPercentage~Family, data = AGF19_Larvae)
summary(mod3)

mod4<-aov(SettledPercentage~CrossPop, data = AGF19_Larvae)
summary(mod4)

mod5<-aov(SettledPercentage~Mom, data = AGF19_Larvae)
summary(mod5)

mod6<-aov(SettledPercentage~Treatment*TimePoint*CrossPop*Mom, data = AGF19_Larvae)
summary(mod6)
################################################
d17 <- subset(AGF19_Larvae, TimePoint == "T17")
d24 <- subset(AGF19_Larvae, TimePoint == "T24")
d40 <- subset(AGF19_Larvae, TimePoint == "T40")
#
s17 <- aov(SettledPercentage~Treatment, data = d17)
summary(s17)
#
s24 <- aov(SettledPercentage~Treatment, data = d24)
summary(s24)
#
s40 <- aov(SettledPercentage~Treatment, data = d40)
summary(s40)
```
