## Supplementary material for "Predicting selection-response gradients of heat tolerance in a widespread reef-building coral": R scripts using in the study: Larvae_settlement_script.R

```
###################################Read in data
library(lattice)
library(plyr)
library(Rmisc)
library(readxl)
#This is my path, you will need to change it to match your local drive path
AGF19_Larvae_T17<-read_excel("D:/PhD/Journal/larvae/sttl_T17_changed.xlsx")
AGF19_Larvae_T24<-read_excel("D:/PhD/Journal/larvae/sttl_T24_changed.xlsx")
AGF19_Larvae_T40<-read_excel("D:/PhD/Journal/larvae/sttl_T40_changed.xlsx")

library(dplyr)
###################################################
#plot with mean point # and errorbar
med17 <- ddply(AGF19_Larvae_T17, .(CrossPop, Temperature), summarise, med = median(SettledPercentage))
#
library(ggplot2)
library(ggpubr)
#
sttl_bp_17<-ggplot(AGF19_Larvae_T17, aes(x=Temperature, y=SettledPercentage))+
  geom_boxplot(aes(color=Temperature),show.legend = FALSE)+ facet_grid(.~CrossPop)+ylim(0,100)+
  scale_color_manual(values = c("35.5C"="coral","27.5C"="skyblue"))+
  ylab("Percentage of settlement")+ggtitle("A") +
  #stat_summary(fun = mean, geom = "point", col = "red") +
  #stat_summary(fun.data = mean_se, geom = "errorbar",col = "red")+
  #theme_bw() +
  #theme(strip.background =element_rect(fill="red"))+
  theme(plot.title = element_text(size = 14, family = "Tahoma", face = "bold"),
        text = element_text(size = 10, family = "Tahoma"),
        axis.title = element_text(face="bold", size = 12),
        axis.text.x=element_text(size = 10))+
  geom_text(data = med17, aes(x = Temperature, y = med, label = med), 
            size = 2, vjust = -1.5)+ stat_compare_means(aes(label = ..p.format..), size = 2.5)
#
#
#
med24 <- ddply(AGF19_Larvae_T24, .(CrossPop, Temperature), summarise, med = median(SettledPercentage))

sttl_bp_24<-ggplot(AGF19_Larvae_T24, aes(x=Temperature, y=SettledPercentage))+
  geom_boxplot(aes(color=Temperature),show.legend = FALSE)+ facet_grid(.~CrossPop)+ylim(0,100)+
  scale_color_manual(values = c("35.5C"="coral","27.5C"="skyblue"))+
  ylab("Percentage of settlement")+ggtitle("B") +
  #stat_summary(fun = mean, geom = "point", col = "red") +
  #stat_summary(fun.data = mean_se, geom = "errorbar",col = "red")+
  #theme_bw() +
  #theme(strip.background =element_rect(fill="red"))+
  theme(plot.title = element_text(size = 14, family = "Tahoma", face = "bold"),
        text = element_text(size = 10, family = "Tahoma"),
        axis.title = element_text(face="bold", size = 12),
        axis.text.x=element_text(size = 10))+
  geom_text(data = med24, aes(x = Temperature, y = med, label = med), 
            size = 2, vjust = -1.5)+ stat_compare_means(aes(label = ..p.format..), size = 2.5)
#
#
#
med40<- ddply(AGF19_Larvae_T40, .(CrossPop, Temperature), summarise, med = median(SettledPercentage))

sttl_bp_40<-ggplot(AGF19_Larvae_T40, aes(x=Temperature, y=SettledPercentage))+
  geom_boxplot(aes(color=Temperature),show.legend = FALSE)+ facet_grid(.~CrossPop)+ylim(0,100)+
  scale_color_manual(values = c("35.5C"="coral","27.5C"="skyblue"))+
  ylab("Percentage of settlement")+ggtitle("C") +
  #stat_summary(fun = mean, geom = "point", col = "red") +
  #stat_summary(fun.data = mean_se, geom = "errorbar",col = "red")+
  #theme_bw() +
  #theme(strip.background =element_rect(fill="red"))+
  theme(plot.title = element_text(size = 14, family = "Tahoma", face = "bold"),
        text = element_text(size = 10, family = "Tahoma"),
        axis.title = element_text(face="bold", size = 12),
        axis.text.x=element_text(size = 10))+
  geom_text(data = med40, aes(x = Temperature, y = med, label = med), 
            size = 2, vjust = -1.5)+ stat_compare_means(aes(label = ..p.format..), size = 2.5)

#####################################3
ggarrange(sttl_bp_17 ,sttl_bp_24, sttl_bp_40,
          common.legend = FALSE, legend = "bottom", ncol = 1)
```
