## Supplementary material for "Predicting selection-response gradients of heat tolerance in a widespread reef-building coral": R scripts using in the study: Larvae_survival_script_finaldata.R

```
rm(list = ls()) 
setwd("C:/Users/amacadam/Dropbox/Kondo Klub/Figures")

library(dplyr)
library(ggplot2)
library(MCMCglmm)
library(nlme)
library(lme4)
library(plotrix)
library(ggpubr)
library(piecewiseSEM)
library(sjPlot)
library(multcomp) #ghlt
library(lsmeans)
library(effects)
library(plotMCMC)
library(MCMCpack)
library(lattice)
library(Rmisc)
library(readxl)
library(rstatix)


data<- read.csv("C:/Users/amacadam/Dropbox/Kondo Klub/Spreadsheets/2019_Larvae_data_curated_JC_final.csv",header=TRUE)
shapiro.test(data$Percent_Survival_atend)

stat.test <- data %>%
  group_by(Cross2) %>%
  wilcox_test(Percent_Survival_atend ~ Temp) %>%
  #adjust_pvalue(method = "bonferroni") %>%
  add_significance()
stat.test 

survival_larv<-ggplot(data, aes(x=Treatment, y=Percent_Survival_atend))+
  geom_boxplot(aes(color=Treatment),show.legend = FALSE)+
  facet_grid(.~Cross2)+ylim(0,100)+
  scale_color_manual(values = c("Hot"="#fb6703", "Ambient" = "#00abff"))+
  theme_grey() +
  theme(plot.title = element_text(size = 14, family = "Tahoma", face = "bold"),
        text = element_text(size = 14, family = "Tahoma"),
        axis.title = element_text(size=14),
        axis.text.x=element_text(size = 14, angle = +45))+
  ylab("Percentage survival at end (%)")+
 theme(strip.background = element_rect(color='black', size=1.5, linetype="solid")) +
  stat_compare_means(label = "p")
survival_larv

#OR
survival_larv + stat_pvalue_manual(stat.test, hide.ns = FALSE, label = "{p}")+
  scale_y_continuous(expand = expansion(mult = c(0.05, 0.1)))

ggarrange(check19, survival_larv, nrow = 2, labels = c("A","B"))

library(png)
library(cowplot)
library(patchwork)
z<-ggdraw()+
  draw_plot(check19, 0, .5, 1, .5)+
  draw_plot(survival_larv, 0, 0, 1, .5)
z

medians<- data %>% group_by(Temp, Cross) %>% dplyr::summarise(Median = median(Percent_Survival_atend, na.rm=T))
write.csv(medians, "C:/Users/amacadam/Desktop/data.csv", row.names=T)
```
