## Supplementary material for "Predicting selection-response gradients of heat tolerance in a widespread reef-building coral": R scripts using in the study: plot and statistical analysis walkthrough.docx

Mean monthly temperature plot (Fig. 1B)

Mean monthly temperature of three sites was plot (see R script AGF19_temperature_plot.R, dataset: Sat_eReefs_AGF19Adult_temps.csv).

Ramping time plot (Fig. 1E)

Ramping time plot showed the experimental temperature and length for adult and larvae (see R script: ramp_figure_AGF18.R, dataset: ramp_figure_AGF18.csv).

Larvae survival plot in cumulative hours (Fig. 2A)

Percentage of larvae survival over time of the experiment was plot in cumulative hours (see R script: Larvae_cumulative_hours.R, dataset: 2019_Larvae_data_curated_JC.csv).

Larvae survival plot and Wilcoxon's test (Fig. 2B)

The mean values of larvae survival at the final time point was compared between each cross at the two treatments (see R script: Larvae_survival_script_finaldata.R, dataset: 2019_Larvae_data_curated_JC_final.csv).

Effects of factors on larval settlement.

Using the dataset Larvae_AGF19_settlement.csv, the analysis of variance of factors (Treatment, TimePoint, CrossPop, Mom) on larval settlement was conducted (see R script Larvae_settlement_factor_script.R)

Larval settlement plot and Wilcoxon's test (Fig. 3)

Comparisons of medians of larval settlement between each cross at the two treatments was conducted separately in each time point (dataset: sttl_T17_changed.xlsx, sttl_T24_changed.xlsx, sttl_T40_changed.xlsx). The Wilcoxon's test was used (see the R script: Larvae_settlement_script.R).

Adult median plot and Wilcoxon's test (Fig. 4A-D)

Four adult responses (bleaching score, ΔF/FM’, percentage of necrosis, and survival) were displayed as boxplots which also showing the Wilcoxon's test (see R script: Adult_median_plot.R, dataset: Adult_AGF19_Tfinal_response.csv).

Adult response model analysis

Bleaching score was analyzed with non-binomial generalized linear model, ΔF/FM’ and percentage of necrosis with linear model, and survival with generalized linear model (see R script: Adult_response_plot.R, dataset: Adult_AGF19_Tfinal_response.csv).

The Percentage changes of adult responses by latitude (Fig. 4E)

The percentage changes of bleaching score, ΔF/FM’, percentage of necrosis, and survival was calculated and put in Adult_latitude_change_result.csv. This data was then used to create line plot (see R script: Adult_latitude_change_plot.R).

Larvae and adult survival density plot (Fig. 5)

The survival percentage of larvae data (see dataset: 2019_Larvae_data_curated_JC_final.csv) was used to create density plot (see R script: Larvae_density_plot.R). The same plot was also done for adult data (see R script: Adult_density_plot.R, dataset: Adult_AGF19_Tfinal_response.csv).

Contour plot of response gradients to selection (Fig. 6)

The contour plot of response to selection (R) for selective potentials (S) and narrow-sense heritability (h^2^) was made (see R script: Survival_contour_plot_KQ.R, dataset: ResponseSelection_final.csv).
