## Supplementary material for "Predicting selection-response gradients of heat tolerance in a widespread reef-building coral": R scripts using in the study: ramp_figure_AGF18.R

```
rm(list = ls()) 
setwd("C:/Users/amacadam/Dropbox/Kondo Klub/Figures")
library(ggplot2)
library(ggpubr)
library(dplyr)
library(forcats)
library(tidyverse)
library(tidyr)
library(plotrix)
library(rstatix)
library(broom)
library(ggthemes)
library(gridExtra)
library(RColorBrewer)

data <- read.csv("C:/Users/amacadam/Dropbox/Kondo Klub/Figures/ramp_figure_AGF18.csv",header=TRUE, fileEncoding="UTF-8-BOM")

plot<-ggplot(data, aes(x=Hours, y=Degrees_C, color=Lifestage))+
  geom_line(size=1)+
 theme_classic2()+
  scale_color_brewer(palette = "Dark2")+
  theme(panel.grid.major.x = element_line(size=12),
        panel.grid.minor.x = element_blank())+
  scale_x_continuous(breaks = c(0, 24,48,72,96, 120, 144, 168, 192, 216, 240, 264, 288, 312))
  scale_x_continuous(minor_breaks = seq(0,312,0.5))
plot
```
