## Supplementary material for "Predicting selection-response gradients of heat tolerance in a widespread reef-building coral": R scripts using in the study: Survival_contour_plot_KQ.R

```
df_sample <-read.csv("D:/PhD/Journal/ResponseSelection_final.csv")
#
library(plotly)
library(dplyr)

df_sample2_hot<-df_sample2 %>% filter(Temperature %in% c("35.5C")) #%>% filter(R.z >0)

density_mom<-ggplot(df_sample2_hot, aes(x = S.x, y = h2.y, fill = R.z, color=Mom))+
  geom_density_2d() + facet_grid(Mom~., scales = "free_y")+scale_color_manual(values = c("#AC9052", "black", "#692F28"))

p<-ggplot(df_sample2_hot, aes(x = S.x, y = h2.y, colour = R.z)) +
  geom_point(aes(colour = R.z))  + facet_grid(Mom~., scales = "free") +
  scale_colour_gradient(low = "#87CCE9", high = "#D76127")

library(patchwork)
density_mom+p

#
```
