## Supplemental tables for "Predicting selection-response gradients of heat tolerance in a widespread reef-building coral"

**Supplementary Material**

**Supplementary Table 1:** Prior to spawning, each *A. tenuis* colony was given an identification number.

| Adult colony | Identification number | Spawning |
| --- | --- | --- |
| Davies 9 | D9 | N |
| Davies 10 | D10 | Y |
| Davies 11 | D11 | Y |
| Davies 12 | D12 | Y |
| Davies 13 | D13 | N |
| Davies 14 | D14 | N |
| Davies 15 | D15 | N |
| Davies 16 | D16 | Y |
| Davies 17 | D17 | N |
| Davies 18 | D18 | N |
| Davies 19 | D19 | N |
| Davies 20 | D20 | Y |
| Esk 1 | E1 | Y |
| Esk 2 | E2 | Y |
| Esk 3 | E3 | Y |
| Esk 4 | E4 | Y |
| Esk 5 | E5 | Y |
| Esk Unknown | EUN | N |
| Keppels 1 | K1 | Y |
| Keppels 2 | K2 | N |
| Keppels 3 | K3 | Y |
| Keppels 4 | K4 | Y |
| Keppels 5 | K5 | N |
| Keppels 6 | K6 | Y |
| Keppels 7 | K7 | Y |
| Keppels 8 | K8 | Y |
| Keppels 9 | K9 | Y |

**Supplementary Table 2:** The genetic crosses of the *Acropora tenuis* larvae used in the settlement assays. Larvae were produced from gametes released during the coral mass spawning event of November 2019. Parent corals were collected from three sites on the GBR – Esk reef (E), Davies reef (D), Keppels reef (K). Multiple cultures were made for a few families, these replicates were included in the settlement assays.

| Family ID | Dam Colony ID | Sire Colony ID | Genetic Cross | Larval Heat Stress | Settlement |
| --- | --- | --- | --- | --- | --- |
| F19 | D10 | E5 | D10 x E5 | N | Y |
| F19 | D10 | E5 | D10 x E5 | Y | Y |
| F35 | D10 | E4 | D10 x E4 | N | Y |
| F35 | D10 | E4 | D10 x E4 | Y | Y |
| F20 | D11 | E5 | D11 x E5 | N | Y |
| F20 | D11 | E5 | D11 x E5 | Y | N |
| F31 | D11 | E3 | D11 x E3 | N | Y |
| F31 | D11 | E3 | D11 x E3 | Y | Y |
| F39 | D11 | K6 | D11 x K6 | N | Y |
| F39 | D11 | K6 | D11 x K6 | Y | N |
| F41 | D11 | K3 | D11 x K3 | N | Y |
| F41 | D11 | K3 | D11 x K3 | Y | N |
| F1 | D12 | D16 | D12 x D16 | N | Y |
| F1 | D12 | D16 | D12 x D16 | Y | N |
| F24 | D12 | E4 | D12 x E4 | N | Y |
| F24 | D12 | E4 | D12 x E4 | Y | Y |
| F29 | D12 | E3 | D12 x E3 | N | Y |
| F29 | D12 | E3 | D12 x E3 | Y | Y |
| F17 | D16 | E5 | D16 x E5 | N | Y |
| F17 | D16 | E5 | D16 x E5 | Y | Y |
| F23 | D16 | E4 | D16 x E4 | N | Y |
| F23 | D16 | E4 | D16 x E4 | Y | N |
| F28 | D16 | E3 | D16 x E3 | N | Y |
| F28 | D16 | E3 | D16 x E3 | Y | Y |
| F36 | D16 | K6 | D16 x K6 | N | Y |
| F36 | D16 | K6 | D16 x K6 | Y | N |
| F37 | D16 | K9 | D16 x K9 | N | Y |
| F37 | D16 | K9 | D16 x K9 | Y | N |
| F38 | D16 | K3 | D16 x K3 | N | Y |
| F38 | D16 | K3 | D16 x K3 | Y | Y |
| F2 | D20 | D16 | D20 x D16 | N | Y |
| F2 | D20 | D16 | D20 x D16 | Y | N |
| F44 | D20 | K3 | D20 x K3 | N | Y |
| F44 | D20 | K3 | D20 x K3 | Y | Y |
| F7 | D20 | D12 | D20 x D12 | N | Y |
| F7 | D20 | D12 | D20 x D12 | Y | Y |
| F32 | D5 | E3 | E5 x E3 | N | Y |
| F100 | E1 | K1 | E1 x K1 | N | Y |
| F100 | E1 | K1 | E1 x K1 | Y | N |
| F101 | E1 | K7 | E1 x K7 | N | Y |
| F101 | E1 | K7 | E1 x K7 | Y | N |
| F98 | E1 | K8 | E1 x K8 | N | Y |
| F98 | E1 | K8 | E1 x K8 | Y | Y |
| F83 | E2 | K8 | E2 x K8 | N | Y |
| F83 | E2 | K8 | E2 x K8 | Y | Y |
| F87 | E2 | K4 | E2 x K4 | N | Y |
| F87 | E2 | K4 | E2 x K4 | Y | Y |
| F10 | E3 | D12 | E3 x D12 | N | Y |
| F10 | E3 | D12 | E3 x D12 | Y | Y |
| F13 | E3 | D10 | E3 x D10 | N | Y |
| F13 | E3 | D10 | E3 x D10 | Y | N |
| F16 | E3 | D11 | E3 x D11 | N | Y |
| F16 | E3 | D11 | E3 x D11 | Y | N |
| F22 | E3 | E5 | E3 x E5 | N | Y |
| F22 | E3 | E5 | E3 x E5 | Y | N |
| F27 | E3 | E4 | E3 x E4 | N | Y |
| F27 | E3 | E4 | E3 x E4 | Y | Y |
| F5 | E3 | D16 | E3 x D16 | N | Y |
| F5 | E3 | D16 | E3 x D16 | Y | N |
| F73 | E3 | E4 | E3 x E4 | N | Y |
| F73 | E3 | E4 | E3 x E4 | Y | Y |
| F74 | E3 | K6 | E3 x K6 | N | Y |
| F74 | E3 | K6 | E3 x K6 | Y | Y |
| F94 | E3 | K9 | E3 x K9 | N | Y |
| F94 | E3 | K9 | E3 x K9 | Y | Y |
| F12 | E4 | D10 | E4 x D10 | N | Y |
| F12 | E4 | D10 | E4 x D10 | Y | Y |
| F15 | E4 | D11 | E4 x D11 | N | Y |
| F15 | E4 | D11 | E4 x D11 | Y | N |
| F21 | E4 | E5 | E4 x E5 | N | Y |
| F21 | E4 | E5 | E4 x E5 | Y | N |
| F33 | E4 | E3 | E4 x E3 | N | Y |
| F33 | E4 | E3 | E4 x E3 | Y | N |
| F4 | E4 | D16 | E4 x D16 | N | Y |
| F4 | E4 | D16 | E4 x D16 | Y | Y |
| F67 | E4 | K6 | E4 x K6 | N | Y |
| F67 | E4 | K6 | E4 x K6 | Y | Y |
| F9 | E4 | D12 | E4 x D12 | N | Y |
| F9 | E4 | D12 | E4 x D12 | Y | N |
| F90 | E4 | K1 | E4 x K1 | N | Y |
| F90 | E4 | K1 | E4 x K1 | Y | Y |
| F11 | E5 | D10 | E5 x D10 | N | Y |
| F11 | E5 | D10 | E5 x D10 | Y | N |
| F14 | E5 | D11 | E5 x D11 | N | Y |
| F14 | E5 | D11 | E5 x D11 | Y | Y |
| F26 | E5 | E4 | E5 x E4 | N | Y |
| F26 | E5 | E4 | E5 x E4 | Y | N |
| F3 | E5 | D16 | E5 x D16 | N | Y |
| F3 | E5 | D16 | E5 x D16 | Y | N |
| F32 | E5 | E3 | E5 x E3 | Y | N |
| F8 | E5 | D12 | E5 x D12 | N | Y |
| F8 | E5 | D12 | E5 x D12 | Y | N |
| F114 | K1 | E1 | K1 x E1 | N | Y |
| F114 | K1 | E2 | K1 x E2 | Y | N |
| F114 | K1 | E3 | K1 x E3 | Y | N |
| F114 | K1 | E4 | K1 x E4 | Y | N |
| F81 | K1 | K7 | K1 x K7 | N | Y |
| F81 | K1 | K7 | K1 x K7 | Y | Y |
| F82 | K1 | K6 | K1 x K6 | N | Y |
| F82 | K1 | K6 | K1 x K6 | Y | N |
| F60 | K3 | D20 | K3 x D20 | N | Y |
| F60 | K3 | D20 | K3 x D20 | Y | N |
| F65 | K3 | K6 | K3 x K6 | N | Y |
| F65 | K3 | K6 | K3 x K6 | Y | N |
| F122 | K4 | E1 | K4 x E1 | N | Y |
| F122 | K4 | E1 | K4 x E1 | Y | N |
| F45 | K6 | D20 | K6 x D20 | N | Y |
| F45 | K6 | D20 | K6 x D20 | Y | Y |
| F47 | K6 | D16 | K6 x D16 | N | Y |
| F47 | K6 | D16 | K6 x D16 | Y | N |
| F48 | K6 | E4 | K6 x E4 | N | Y |
| F48 | K6 | E4 | K6 x E4 | Y | N |
| F49 | K6 | E2 | K6 x E2 | N | Y |
| F49 | K6 | E2 | K6 x E2 | Y | N |
| F51 | K6 | K9 | K6 x K9 | N | Y |
| F51 | K6 | K9 | K6 x K9 | Y | Y |
| F52 | K6 | K3 | K6 x K3 | N | Y |
| F52 | K6 | K3 | K6 x K3 | Y | N |
| F116 | K7 | E4 | K7 x E4 | N | Y |
| F116 | K7 | E4 | K7 x E4 | Y | N |
| F80 | K7 | K6 | K7 x K6 | N | Y |
| F80 | K7 | K6 | K7 x K6 | Y | N |
| F103 | K8 | K8 | K8 x E2 | N | Y |
| F103 | K8 | E2 | K8 x E2 | Y | Y |
| F106 | K8 | E1 | K8 x E1 | N | Y |
| F106 | K8 | E1 | K8 x E1 | Y | N |
| F53 | K9 | D20 | K9 x D20 | N | Y |
| F53 | K9 | D20 | K9 x D20 | Y | N |
| F54 | K9 | D11 | K9 x D11 | N | Y |
| F54 | K9 | D11 | K9 x D11 | Y | N |
| F55 | K9 | D16 | K9 x D16 | N | Y |
| F55 | K9 | D16 | K9 x D16 | Y | N |
| F57 | K9 | E3 | K9 x E3 | N | Y |
| F57 | K9 | E3 | K9 x E3 | Y | N |
| F58 | K9 | K6 | K9 x K6 | N | Y |
| F58 | K9 | K6 | K9 x K6 | Y | N |
| F59 | K9 | E3 | K9 x E3 | N | Y |
| F59 | K9 | E3 | K9 x E3 | Y | Y |

**Supplementary Table 3:** Statistical analyses of random effects of experimental tank and fragment stick on the physiological responses of adult fragments

| Adult response | Random effect analysis (P Values) | |
| --- | --- | --- |
|  | Tank | Stick |
| Bleaching | 0.2953 | 0.1087 |
| Effective quantum yield | 0.4195 | 0.075 |
| Percentage of necrosis | 0.3358 | 0.118 |
| Survival | 0.304 | 0.111 |

**Supplementary Table 4**: Wilcoxon's test of larval settlement comparing between heat and control treatment at 17 h, 24 h and 48 h

| Population cross | 17 hrs | 24 hrs | 48 hrs |
| --- | --- | --- | --- |
| DxD | 0.00021 | 0.00016 | NA (all larvae died) |
| DxE | 1.2e-06 | 4.3e-10 | 9.9e-09* |
| DxK | 2.2e-06 | 3.2e-09* | 1.1e-07 |
| ExD | 1.8e-07 | 2.7e-11 | 2.1e-09 |
| ExE | 1.7e-06 | 4.3e-08 | 1.2e-07 |
| ExK | 1.4e-07 | 5.3e-13 | 3e-12 |
| KxD | 1.1e-06 | 1.7e-07 | 2.8e-06 |
| KxE | 1.4e-07 | 2.2e-10 | 1.1e-07 |
| KxK | 5.8e-06 | 3.2e-09 | 1.5e-07 |

**Supplementary Table 5:** Adult coral colonies used for adult heat stress experiments and for larval family crosses

| Adult ID Number | Used in Adult Heat Stress Experiment | Used for Larval Family Crosses |
| --- | --- | --- |
| D9 | Y | N |
| D10 | N | Y |
| D11 | N | Y |
| D12 | N | Y |
| D13 | Y | N |
| D14 | Y | N |
| D15 | N | N |
| D16 | N | Y |
| D17 | N | N |
| D18 | Y | N |
| D19 | N | N |
| D20 | Y | Y |
| E1 | Y | Y |
| E2 | Y | Y |
| E3 | N | Y |
| E4 | N | Y |
| E5 | N | Y |
| EUN | Y | N |
| K1 | N | Y |
| K2 | N | N |
| K3 | Y | Y |
| K4 | N | Y |
| K5 | N | N |
| K6 | Y | Y |
| K7 | Y | Y |
| K8 | N | Y |
| K9 | N | Y |
