## Supplementary material for "Predicting selection-response gradients of heat tolerance in a widespread reef-building coral": Responses to comments from journal's reviewer

| **Reviewer comment number** | **Section** | **Comment/ Suggestion** | **Response** | **Changes** |
| --- | --- | --- | --- | --- |
| Reviewer  1.1 | Data and Code | Unable to assess manuscript from a technical perspective as the URL for the repository led to a 404 error and unable to find it in the LaserKate repositories GitHub page.  I would recommend that the repository be made public and/or that it be stored on a more permanent repository such as Zenodo or Dryad. | A zipped folder with all the R scripts has been added to the submission system for Reviewers.  All scripts will be uploaded to the Github location at the time of publication. | NA |
| 1.2 | Introduction  Lines 81-88 | The last few sentences of this paragraph seem to jump around from specific effects of temperature on larvae physiologically to how these physiological impacts affect adaptation and back again. It seems like all of this information is important to include and some of it could be expanded upon.  I would therefore recommend that this paragraph be split into two to give each point room to grow, with the first paragraph being about the specific effects of temperature on larvae physiologically and the second relating this information to its impact on adaptation. | This paragraph has been divided and re-organized as suggested. | Lines 88-107:  “Reductions in dispersal distances have the potential to reduce gene flow between populations, thereby potentially accelerating population differentiation and local adaption (Bassim et al., 2002). Additionally, some coral populations rely on the dispersal of larvae from neighbouring populations for the addition of beneficial genes/genotypes to their gene pool (Munday et al., 2009; Quigley et al. 2019). With reduced reef connectivity, populations may not be able to adapt rapidly enough to cope with increasing SSTs (Quigley et al. 2019) or may struggle to regenerate following mass bleaching and mortality (Bassim et al., 2002). There is scope for adaptation of corals to increasing temperatures (Matz et al., 2018). Further, the temperatures at which bleaching is occurring have increased by ~0.5 °C from 1998 to 2017, suggesting an increase in more heat-adapted genotypes within coral populations (Sully et al., 2019), potentially through processes like selective sweeps (Quigley et al., 2019a). Locally, there have been reported increases in coral cover in both the central and southern GBR (AIMS, 2020), with evidence to suggest that warm or “extreme” habitats harbour an increased number of thermally adapted genotypes with the potential to transmit heat tolerance (Quigley et al., 2020a; Schoepf et al., 2019). However, current estimates of the rate of SST increases suggest that these increases may exceed the potential rate of fixation of beneficial genetic variants given factors like currents and reef topology (Quigley et al., 2019a). Additionally, the annual increase in SSTs has extended the period at which ‘summer’ temperatures occur, further increasing global bleaching by reducing potential recovery periods that occur when coral populations are exposed to cooler ‘winter’ temperatures (Heron et al., 2016). Taken together, this suggests that corals adaptive potentials may be constrained.” |
| 1.3 | Introduction  Line 88 | It is unclear here if “This information” is about the physiological responses of  larvae or on dispersal. If it is about the physiological responses, I would just change the wording from “largely” to “relatively” unknown to acknowledge those who have looked into it. I would also recommend citing McLachlan, Price, Solomon & Grottoli (2020), who found that 95% percent of studies focused on adult responses and only 2 and 1% on pre-settled and settlement stages, respectively. Additionally, if this information refers to the physiological responses of early life stages to warming, I suggest giving a very brief 1-2 sentence synthesis of current knowledge of pre-settled and settlement life stages beyond that said above (line 80). If “This information” is instead about dispersal then you can ignore my musings. | This has been clarified and the additional information and reference added. | Lines 70-73:  “Despite this, information on larval physiological responses is relatively unknown for coral early life-history stages (McLachlan et al., 2020), with 95 % of studies focusing on adult responses and only 2 % and 1 % on pre-settled and settlement stages, respectively (McLachlan et al., 2020)” |
| 1.4 | Introduction  Lines 116-118 | Because this journal is read by a broad audience, I would provide brief explanations for some of the more jargon-y terms here that may be unfamiliar to a broader audience, especially, “narrow-sense heritability”, “selection coefficients”, and “breeders equation”. | Extra detail has been added as requested. | Lines 122-130:  “Further, evolutionary models incorporating metrics like narrow-sense heritability (h^2^; the phenotypic traits within an organism that arise from allele inheritance; Evans et al., 2018), selection coefficients (S; relative fitness of a phenotypic trait; van Tienderen and de Jong, 1994), and responses to selection are commonly measured (Falconer and Makcay, 1996), and allow for the prediction of organismal responses to future stressors. Thus far, models incorporating the “breeders equation” (R = h^2^S; used to predict the effect of selection pressures on phenotypic traits; Falconer and Makcay, 1996) have demonstrated the utility of evolutionary modelling for predicting the potential of the coral’s algal symbionts to confer increased survival (Quigley et al., 2018).” |
| 1.5 | Methods  Larval rearing | Can you include more information about larval rearing in the methods or in the supplementary materials? This information is important to include for replication purposes. Specifically, methodology for the control and measurement (both method and frequency) of water flow rate, light intensity, salinity, pH, and especially temperature should be included | This information has been added. | Lines 196-200:  “Once fertilization was confirmed, embryos from each separate bowl were transferred into separate 15 L constant flow-through conical tanks with 0.2 μm FSW in a temperature-controlled room, such that the temperature of each cone was maintained at 27.5 °C, *p*CO_2_ 400±60ppm, ambient light (i.e. non-photosynthetic), and salinity of 35 psu.” |
| 1.6 | Methods  Larval heat stress experiment | Can you provide a brief explanation of how temperatures for heat stress were chosen and why the adults were subjected to lower heat than the larvae? | This has been included. | Lines 255-257:  “Temperatures for larval and adult experiments were chosen in order to compare with previous work performed in the same region using a cross design (Dixon et al. 2015; Quigley et al. 2020b).” |
| 1.7 | Methods  Settlement experiment | Can you explain the criteria for the selection of the subsetting of 69 crosses from the original 85? Why specifically were 69 chosen? | Only 69 of the 85 families of larvae were chosen because the remaining 16 families had an insufficient number of larvae remaining in the stock tanks for experimental use. This has been added to the Methods. | Lines 216-218:  “Following the larval heat stress experiment, only 69 of the original 85 crosses contained sufficient larval stock for further experimental use. These 69 crosses were used to investigate the effect of thermal stress on larval settlement behaviour.” |
| 1.8 | Methods  Adult heat stress experiment | For replication purposes, can you include the same information about adult culturing as requested for larval rearing (above)? | This has been added. | Lines 233-234:  “Colonies were kept in outdoor aquaria under the following conditions before fragmentation: 0.2 μm FSW, 27.5 °C, *p*CO_2_ 400±60 ppm, and salinity of 35 psu.” |
| 1.9 | Methods  Adult heat stress experiment | To clarify, were the colonies used for the adult heat stress experiment the same corals used for spawning? | Due to some corals not spawning and necrosis of adult colonies, not all adult colonies used for the heat stress experiment were used for spawning and vice versa. A table listing which adults were used for the heat stress experiment and for spawning has been added to the supplementary material. | This information is available in Supplementary Table 5. |
| 1.10 | Methods  Adult heat stress experiment | Can you add a brief description of how genotypes for the adult corals were determined? If the genotypes were known a priori, this information could be included in the Coral Colony Collection paragraph. | This information has been added. | Lines 167-169:  “Corals were identified as individual genotypes during collection by removing colonies approximately 10 m apart if possible. Genotypes were not known *a priori*.” |
| 1.11 | Statistical analysis  Larval heat stress experiment | Can you describe how you checked that the statistical assumptions of the wilcox.test were met by your data? | This has been added. | Lines 272-273  “After testing data normality and homogeneity of variance with diagnostic plot using "stats" package (R Core Team, 2021) …” |
| 1.12 | Statistical analysis  Adult heat stress experiment | To clarify (in reference to lines 275-276), were separate models made for each variable (bleaching, photophysical response, percent necrosis, and  survival? | Yes. Each model was chosen to fit those independent responses. For example, survival data were recorded in binary (1 and 0) so it was best to use a binomial generalised linear model. | Lines 300-302:  "Finally, the negative binomial generalized linear model was used for bleaching, linear model for ΔF/FM’ and the percentage of necrosis, and generalized linear model for survival" |
| 1.13 | Statistical analysis  Adult heat stress experiment | Based on the bleaching score assignment described above, it seemed like bleaching score would be a discrete variable (integers ranging 0-6) instead of a continuous variable. Can you also clarify why bleaching was treated as a continuous variable instead of a discrete variable and if any transformations were needed on this dataset? | Bleaching is treated as discrete | Lines 295-296:  “The metric ΔF/FM’ and the percentage of necrosis were treated as continuous variables, and temperature treatment and population were set as fixed factors in each model” |
| 1.14 | Statistical analysis  Adult heat stress experiment | Include the specific type of model used to analyse the effects for each variable in this paragraph. For example, in the results (line 364), it was mentioned that a negative binomial generalized linear model was used for bleaching score, but this is not mentioned in the methods section. | This has been added. | Lines 300-302:  “Finally, the negative binomial generalized linear model was used for bleaching, linear model for ΔF/FM’ and the percentage of necrosis, and generalized linear model for survival.” |
| 1.15 | Discussion  Larval settlement  Paragraph 1  Line 519 | Contextualise your  pre-settlement survivorship results | Please see response in 2.18 | Lines 552-570:  “Previous breeding experiments have demonstrated the transfer of increased offspring survival from parents sourced from warm reefs when reproductively crossed with cooler reefs (Dixon et al., 2015), or at least one parent from warmer reefs (Quigley et al., 2020b), suggesting genetic contribution to offspring. In this study, larval survival was high in the crosses whose mother sourced from either Esk or Davies. However, it is currently unknown whether an increased propensity for settlement at high temperatures is also transferable using colonies sourced from warmer reefs to achieve an enhancement in settlement success. Although settlement is a heritable trait under control conditions (h2 = 0.49; Meyer et al., 2009), the overall heritability is low relative to other fitness-related traits. Moreover, it is well known that settlement in corals is negatively impacted by heat. For example, early life-stage Acropora tenuis settlement decreased by 100% when exposed to +5 °C over ambient temperature (Humanes et al., 2016), and by 55 % when combined with suspended sediments treatment (Humanes et al., 2017). Diploria strigose larvae demonstrated a decrease in the settlement behaviour at temperatures exceeding 30 °C compared to control (Bassim and Sammarco, 2003) and Acropora palmata settlement decreased by 25 % at 31.5 °C compared to 28 °C (Randall and Szmant, 2009). The lack of strong differences in settlement success between the crosses here may be reflected in the roughly equal heat tolerance of both Davies and Keppels corals, suggesting that both populations are roughly equivalent in tolerance and therefore did not produce strong differences in settlement of larvae. Combined, these previous results suggest that selection should act on this important trait over time if oceans continue to warm.” |
| 1.16 | Discussion  Larval settlement  Paragraph 2 | A couple of sentences at the beginning of this paragraph reminding the reader of your settlement results would help to contextualise this part of the discussion with your study. | A sentence summarising the settlement results has been added to the start of this paragraph. | Lines 572-573:    “The results from this study show that, when exposed to heat, larvae from all crosses significantly decreased in their settlement behaviour relative to the control temperature. During periods of…” |
| 1.17 | Discussion  Larval settlement  Paragraph 3 | Here you discuss how in the crosses ExK and DxE the maternal corals were from the central warmer reefs. Is there a reason to believe that the  maternal coral would have more of an effect on settlement than the parental coral (seeing that Keppel provided the sperm for one of these more successful crosses)? | Information explaining the maternal effect on heat tolerance has been added. | Lines 593-595:  “Previous research suggests mitochondrial DNA (mtDNA) plays a large role in the thermal resistance of corals, alluding to a high maternal effect on the heat tolerance of coral offspring (Dixon et al., 2015; Quigley et al. 2020).” |
| 1.18 | Discussion  Larval settlement  Paragraph 3 | It also seems to be important to discuss these preliminary findings in the context of the success of the adult Keppels corals discussed earlier. | Further information has been added. | Lines 565-568:  “The lack of strong differences in settlement success between the crosses here may be reflected in the roughly equal heat tolerance of both Davies and Keppels corals, suggesting that both populations are roughly equivalent in tolerance and therefore did not produce strong differences in settlement of larvae.” |
| 1.19 | Abstract  Line 32 | I could be mistaken, but I think the word introgression specifically refers to crosses between different species whereas here the authors are specifically referring to crosses between populations of the same species. Maybe change the word introgression to outcrossing here? | This has been changed as suggested. | Lines 29-31:  “…to assess acquired heat tolerance via outcrossing of offspring phenotypes by comparing five physiological responses (photosynthetic yields, bleaching, necrosis, settlement, and survival).” |
| 1.20 | Abstract  Line 32 | The word multiple here seems to indicate a larger number than three. For clarification purposes, I would just replace “multiple” with “three”. | This is a very good point for clarity. We have replaced both uses of “multiple” with actual numbers. | Lines 29-32:  “…offspring phenotypes by comparing **five** physiological responses (photosynthetic yields, bleaching, necrosis, settlement, and survival). Adaptive potentials and physiological reaction norms were calculated across **three** life-stages to integrate heat tolerance at different biological scales.” |
| 1.21 | Abstract  Line 34-35 | The wording of the sentence “Adult responses at heat… compared to the cooler reef” is a little confusing. After reading the manuscript I think it means that the adult responses were greater than the larval responses? If so, this could be clarified by adding “than” so that it reads “Adult responses at heat were similar but greater than in larvae”. | This has been clarified. | Lines 34-35:  “At heat, adults were less variable compared to larval responses in warmer reefs compared to the cooler reef.” |
| 1.22 | Introduction  Line 80 | It seems like something is missing from the sentence “Increased SSTs may… reducing pre-competency periods”. If not, I would just replace the comma before “reducing pre-competency periods” with and, which, or thereby or something to that effect. | “Thereby” has been added to the sentence. | Lines 81-83:  “Increased SSTs may induce premature metamorphosis by increasing larval metabolic activity and development rates, thereby reducing pre-competency periods (Heyward and Negri, 2010).” |
| 1.23 | Introduction  Line 81-84 | The sentence starting on line 81 with “This reduction” is a bit of a run-on sentence which I think dilutes its contribution to the paragraph and makes it a little confusing to read. If you can split it into two sentences I think this would improve the readability of this line and add a little punch to the paragraph | Good suggestion. This has been edited and split into two sentences, with the latter starting a new paragraph. | Lines 83-89:  “This reduction may influence subsequent larval dispersal distances and result in settlement occurring in suboptimal conditions (Edmunds et al., 2001). Therefore, it is important to incorporate these responses into predictive models of ecosystem change given their flow-on effects into key demographic and population level dynamics.  Reductions in dispersal distances have the potential to reduce gene flow between populations, thereby potentially accelerating population differentiation and local adaption (Bassim et al., 2002).” |
| 1.24 | Introduction  Line 122 | Add “of populations” or another subject to clarify the sentence. For example, “to harness the natural adaptive potential of coral populations for conservation and restoration”. And/or changing the word “natural” to “existing” can have a similar effect, such as “to harness the existing adaptive potential of coral reefs for conservation and restoration.” | The sentence has been changed accordingly to add clarity. | Lines 130-132  “Similarly, these principles can be applied to the coral host to quantify adaptive potentials, in which this information can then be used to harness the **existing** adaptive potential **of coral populations** for conservation and restoration.” |
| 1.25 | Methods  Lines 158-160 | The sentence starting with “Colonies from each location were acclimated…” may fit better contextually at the end of this paragraph or the beginning of the next paragraph. | Good suggestion, this has been added. | This sentence has been moved to the end of the paragraph (materials and methods -> coral colony collections -> paragraph 1).  Lines 173-174:  “Colonies from each location were acclimated in separate outdoor holding tanks under constant 0.2 μm filtered seawater (FSW) flow-through conditions, maintained at 27.5 °C.” |
| 1.26 | Methods  Lines 186-190 | This sentence about population-level crosses may fit better at the end of the previous paragraph. | Good suggestion, this has been added. | This sentence has been moved to the end of the previous paragraph (materials and methods -> coral spawning, selective breeding, and larval rearing  -> paragraph 1).  Lines 188-193:  “Population-level crosses are referred to as follows with the maternal colony first and then paternal colony, in which intrapopulation crosses (filled circles; Fig. 1D) are ExE (EskxEsk), KxK (KeppelsxKeppels), DxD (DaviesxDavies), and interpopulation crosses are ExK (EskxKeppels), ExD (EskxDavies), KxE (KeppelsxEsk), KxD (KeppelsxDavies), DxE (DaviesxEsk), and DxK (DaviesxKeppels) (open circles; Fig. 1D).” |
| 1.27 | Methods  Lines 190-191 | This sentence about none of the crosses perishing may make more sense contextually at the beginning of the next paragraph after saying that thirty larvae were sampled from each cross. | This has been moved as suggested. | This sentence has been moved to the end of the next paragraph (materials and methods -> larval heat stress experiment  -> paragraph 1).  Lines 212-213:  “No crosses perished post-fertilization before sampling took place.” |
| 1.28 | Methods  Lines 233-234 | The part of the sentence, “such that each tank included the same genotypes from each population in all tanks” is a little confusing. Maybe reword it to the effect of “such that each tank included all genotypes from each population”? | This sentence has been changed accordingly. | Line 249:  "…, such that each tank included all genotypes from each population" |
| 1.29 | Methods  Line 238 | Is “Sunrise” in the sentence “Sunrise was set at 21:00 hours” supposed to be “Sunset”? Or was the light cycle flipped so that spawning occurred during the day? | This has been changed. | Lines 254:  "Sunrise was set at 09:00 h." |
| 1.30 | Results  Line 334 | When looking at the settlement rate reported here for the heat stress corals, I was a little confused by the sentence “DxE and ExK showed the greatest percentage in settlement…” and saw the median value was 0.00%. Then I looked at the boxplot and saw the upper quartile was much higher than the other crosses. To clarify this, I would recommend that the upper quartile should also be reported here. | This has been added as suggested | Lines 357-358:  “… in which DaviesxEsk was the only population cross that settled at heat (upper quartile = 100%).” |
| 1.31 | Results  Line 364 | Here I believe “mean” should say “median” based on the methods section. | Thank you for spotting this. This has been changed. | Line 390-391:  “…on the median bleaching score of coral fragments…” |
| 1.32 | Results  Line 436 | Are these values reported after the mean (X.XX ± X.XX%) standard deviation or standard error? I recommend that this be included for just the first occurrence. | It's SE | (Mean±SE) has been added after the first X.XX±X.XX % value in the paragraph ‘adult responses to heat stress at the population level’  Line 392:  “…score of 4.75 ±0.14 (mean ±SE; median = 5.00) whilst…” |
| 1.33 | Results  Line 440 | For the sentence, “When Davies eggs were crossed with sperm from the other central reef, heat tolerance increased by 14.46% compared to DxE”, I was a little confused because I thought DxE was the Davies eggs and Esk (the other central reef?) sperm? | At heat, DxD mean of survival = 55.78 and DxE = 70.24. DxE survival was higher than DxD (purebred) by 14.46% | Lines 467-468:  "When Davies eggs were crossed with sperm from the other central reef, heat tolerance increased by 14.46 % compared to purebred" |
| 1.34 | Discussion  Line 589 | Extra “)” after Quigely et al., 2020b) | This has been changed. | Extra bracket has been removed. |
| 1.35 | Discussion  Line 595 | Change “mechanisms is” to “mechanisms are” | This has been changed. | Lines 648-650:  “Although the underlying mechanisms **are** unknown here, the drivers of bimodality may be linked to biochemical complexity (Rezende and Bozinovic, 2019).” |
| 1.36 | Figure 1 | It looks like the figure legend of Figure 1E compressed during resizing so that it is unclear what the circles and triangles symbolize | This information has been added to the legend. | Lines 919-920:  “Circles indicate the end of each experimental time point, triangles indicate specific points along the experimental timeframe.” |
| 1.37 | Table S2 | It looks like there are four rows for Family 114 and only two rows for every other family. It might supposed to be like that, but I brought it up in case it was an error. | This has been clarified. | Supplementary table 2:  Multiple cultures were made for a few families, these replicates were included in the settlement assays. |
| **Reviewer**  **2.1** | Throughout | Omit reef acronyms - they make the text (especially the results) hard to follow. In figures and/or writing throughout remind the reader why each reef presents a unique opportunity to study the stress endurance of corals. Consider labelling by something that is more tangible to the reader (i.e., temperature regime or reef type) as opposed to acronyms (DxG, GxE). | The acronyms have been changed | This has been changed throughout the text. |
| 2.2**.1** | Experimental Design | It is unclear why the *A. tenuis* are exposed to such harsh temperature treatments  (36C, ~+8C from experiment control, +10-12C reef MMM per L162). | This justification has been added to Methods. | Lines 255-257:  “Temperatures for larval and adult experiments were chosen in order to compare with previous work performed in the same region using a cross design (Dixon et al. 2015; Quigley et al. 2020b).” |
| 2.2.2 | Major concern 2 | I worry whether the ms overinterprets the  survival of few (outlier) juvenile corals (Fig 3, L347, L355) as differential population responses. Can the adaptive capacity of a population be assessed effectively if the majority of individuals perish during thermal challenges? | We had replicate families within each population crosses, so we are averaging their responses. So even though there were only a few larvae from each family, the responses were observed across multiple families as biological replicates (n=85). | Lines 185-188:  “… create 85 distinct coral families with at least one cross per family. These families, observed as biological replicates, comprising of intrapopulation (within the same reef) and interpopulation (between different reefs) crosses …” |
| 2.2.3 |  | How do these settlement rates compare to other studies? | Further comparisons to the literature have been added. | Lines 560-565:  “For example, early life-stage *Acropora tenuis* settlement decreased by 100% when exposed to +5 °C over ambient temperature (Humanes et al., 2016), and by 55 % when combined with suspended sediments treatment (Humanes et al., 2017). *Diploria strigose* larvae demonstrated a decrease in the settlement behaviour at temperatures exceeding 30 °C compared to control (Bassim and Sammarco, 2003) and *Acropora palmata* settlement decreased by 25 % at 31.5 °C compared to 28 °C (Randall and Szmant, 2009)” |
| 2.2.4 |  | How does the biological rationale for the temperature challenge selected follow the recommendations in Grottoli *et al* 2021 Ecol Applications table  1? (I recognize that this ms might not have been available prior to the experimentation in 2019 but it would be  helpful for the ms to comment on how the experimental design compares to other GBR juvenile coral work). | Grottoli et al. provide the comparisons of wide-range coral experiments on bleaching. And one part of it is coral ex-situ condition mimicking the increasing SST. The baseline temperature, given from Grottoli et al, is Maximum Monthly Mean. And the experiment temperature falls up on MMM between +3°C and +9°C (+9°C can be found with heat pulse experiment).  McLachlan et al. also provide variability in coral heat stress experiments. For moderate-term studies, 8-30 days, the experiment temperature was normally set up to 4.3 ± 2.0 °C | Lines 608-613:  “…resembling short-term acute heat stress temperature range (Grottoli et al., 2021; McLachlan et al., 2020). However, the experimental temperature of many studies does not exceed +5 °C above the control temperature (Humanes et al., 2016; Humanes et al., 2017; McLachlan et al., 2020; Quigley et al., 2020b). Hence, our result could reflect the contribution of higher-than-threshold temperature treatment.” |
| 2.3 | Title | Consider replacing “wide-ranging” with widespread or generalist | This has been modified. | Title:  “Predicting selection-response gradients of heat tolerance in a widespread reef-building coral” |
| 2.4 | Abstract  Line 34 | Is “limited improvement” an appropriate way to describe these results? | This has been clarified. | Lines 32-34:  “Selective breeding improved larval survival to heat by 1.5 - 2.5x but did not result in substantial enhancement of settlement, although population crosses were significantly different.” |
| 2.5 | Abstract  Line 36 | Could this also be explained by transport-induced tissue necrosis? | Further explanation has been added to the Discussion. | Lines 533-536:  “Both Davies and Esk corals were collected during the sample trip with the same level of handling. This suggests that transport issues were not the cause of their diminished health state but instead point towards population-level differences between these corals.” |
| 2.6 | Introduction  Line 73 | it is unclear why the *A. tenuis* are exposed to such harsh temperature treatments  (36C, ~+8C from experiment control, +10-12C reef MMM per L162) | This justification has been added to Methods. | Lines 255-257:  “Temperatures for larval and adult experiments were chosen in order to compare with previous work performed in the same region using a cross design (Dixon et al. 2015; Quigley et al. 2020b).” |
| 2.7 | Introduction  Line 110 | Is there considerable population divergence at the subspecies level? | Further information has been added. | Lines 116-120:  “This geographically distinct habitat could contribute to sub-speciation, seen in other marine invertebrates that have different thermal thresholds, like subspecies of *Crassostrea gigas* (Ghaffari et al., 2019). However, it is less clear whether similar patterns in heat tolerance at the individual coral genotype scale to the population level, a trend seen in fish, but not insect or plant species collected across temperature clines (Payne et al., 2021; Rezende and Bozinovic, 2019)” |
| 2.8 | Methods  Line 166 | Were genotypes verified? Were any steps taken to avoid sampling of clones? | This information has been added. | Lines 167-169:  “Corals were identified as individual genotypes during collection by removing colonies approximately 10 m apart if possible. Genotypes were not known *a priori*.” |
| 2.9 | Methods  Line 210 | Include light cycle information | This information was added to the Methods | Lines 223-225:  “... (12:12 day:night light cycle, 170 to 180 PAR, Steridium E-500, lights: Sylvania FHO24W/T5/865 and Innova 4230, light: Sylvania F15W/865)...” |
| 2.10 | Methods  Line 230 | What type of microalgae? | This has been added. | Lines 245-246:  “…microalgae (*Parmelia sulcate*, T-ISO, *Chaetoceros muelleri, Nannochloropsis oceania*, and *Dunnaiella sp*).” |
| 2.11 | Methods  Line 247 | Why not present PSII photochemical efficiency data (Grotolli *et al* 2021)? It is attainable from a diving PAM and presents data frequently reported in studies (McClanahan *et al* 2020 Coral reefs) and therefore makes the stress evaluations in this study more comparable | Further explanation has been added. | Lines 265-266:  “Effective quantum yield **of photosystem II** (ΔF/FM’), which is the efficiency of photon absorption, was measured using the Diving PAM (Walz, 2018)” |
| 2.12 | Methods  Line 293 | Include Breeder's equation | This has been added. | Line 314:  “The breeder’s equation, R = h^2^S (Falconer and Makcay, 1996)…” |
| 2.13 | Results | Very difficult to follow (see major concern 1). Consider reorganising results subheadings to clearly distinguish which sets of results refer to which sets of experiments. | This is a good point. We have included additional subheadings to improve clarity. | Please see additional subheadings provided in the Results section. |
| 2.14 | Results  Section starting at line 416 | Unclear why the selection gradient results are not described with the other larval results? | The selection gradient results incorporate both larval and adult responses. We therefore felt they were best described last after the initial results on each life-stage are described first. | NA |
| 2.15 | Discussion  Line 487 | Incorporate additional references - Parkinson et al 2016 reported widespread gene expression differences of Breviolum cultures at the within- and between-species levels. No temperature stress experiment was conducted | We have replaced this reference (Berkelmans and van Oppen, 2006) with a new reference.  The work of Parkinson et al. (2016) gave an insight of inter-species (four species of Symbiodinium spp. in clade B) and inter-species (strains). It showed different photosynthesis-related genes content among different species (subclade level) and some gene difference between strains (subspecies level) that could affect photosynthesis performance.  For clade level of variation response to heat can be found in many other studies such as bay et al., 2016 (https://doi.org/10.1098/rsos.160322), Berkelmans & van Oppen, 2006 (https://doi.org/10.1098/rspb.2006.3567) | Lines 513-516:  “This variation may be underpinned by genetic variation or by responses mediated by non-genetic mechanisms, like changes in the microbiome (e.g. bacteria or Symbiodiniaceae), in which algal symbiont assemblages may shape corals’ responses to heat stress **(Berkelmans and van Oppen, 2006)**.” |
| 2.16 | Discussion  Paragraph starting at line 491 | Can these results be incorporated in the broader literature? How do the results from this study compare to other studies conducted on these reefs? *A. tenuis*? | Further information has been added. | Lines 523-527:  “*Acropora tenuis* in both central and southern reefs generally hosts dominant abundances of *Cladocopium* (Rocker et al., 2017; Ulstrup and van Oppen, 2003), which could contribute to the similarity in their physiological performance, whereas symbionts from Davies reef or the host corals themselves may have lower initial tolerances but are able to recover and survive equally well.” |
| 2.17 | Discussion  Line 500 | Were necrosis issues due to transport issues? | Further information has been added here. | Lines 533-536:  “Both Davies and Esk corals were collected during the sample trip with the same level of handling. This suggests that transport issues were not the cause of their diminished health state but instead point towards population-level differences between these corals.” |
| 2.18 | Discussion  Line 519 | Interesting! I think the ms could be improvised by further elaborating on the impact of thermal history on juvenile coral physiology and incorporating more references from this area | Further information has been added. | Lines 552-556:  “Previous breeding experiments have demonstrated the transfer of increased offspring survival from parents sourced from warm reefs when reproductively crossed with cooler reefs (Dixon et al., 2015), or at least one parent from warmer reefs (Quigley et al., 2020b), suggesting genetic contribution to offspring. In this study, larval survival was high in the crosses whose mother sourced from either Esk or Davies.” |
| 2.19 | Discussion  Line 561 | Agreed. Are there other studies that compare acute and moderate thermal challenges to juvenile coral settlement that could help explain these patterns? | Please see explanation and changes in 2.2.4 | Lines 608-613:  “…resembling short-term acute heat stress temperature range (Grottoli et al., 2021; McLachlan et al., 2020). However, the experimental temperature of many studies does not exceed +5 °C above the control temperature (Humanes et al., 2016; Humanes et al., 2017; McLachlan et al., 2020; Quigley et al., 2020b). Hence, our result could reflect the contribution of higher-than-threshold temperature treatment.” |
| 2.20 | Discussion  Line 588 | What is the theoretical range of H2 and what does the max/min mean? | This information has been added. | Lines 638-640:  “Narrow-sense heritability ranges from 0 – 1, where 0 is indicative of no genetic contribution to trait variance and 1 is complete dominance of genetics in determining trait variance (Falconer and Makcay, 1996).” |
| 2.21 | Discussion  Line 601-604 | 1. I appreciate the lengths the ms took to include the selection differentials for the 3 sites. However, this section is also difficult to contextualise and citations are sparse.  2. How do these values compare to a theoretical population with/without adaptive potential?  3. Are there examples of other marine invertebrates that can be incorporated? | This information has been added. | 1. Further references have been added in each point below:  2. Lines 660-662:  “In comparison, responses to selection in well-studied systems like aquaculture species, including fish and shellfish, averaged about ~13 % for growth and 4.9 % for survival (Gjedrem and Rye, 2018).”  3. In addition to the example on mollusks that was already incorporated, we have provided further comparisons:  Lines 666-669:  “In this case, mollusks reported a 1.7 - 2.1x difference in responses between larvae and adults, which mirrors reports seen in other invertebrates like brine shrimp (2.7 - 4.9x; Norouzitallab et al., 2014), copepod (Tangwancharoen, 2014), and other invertebrates (Pandori and Sorte, 2019).” |
| 2.22 | Figure 1A | The tan, black and maroon icons are difficult to see over the grey map | This has been edited for clarity by adding a white circle inside the points. | See New Figure 1A |
| 2.23 | Figure 2A | Difficult to distinguish tan, black and maroon lines over grey panel | This colour has been darkened for clarity. | See New Figure 2A |
| 2.24 | Figure 5B-D and Figure 3 | Are these presentations of the same survivorship data? I think the presentation in Fig 3 is a more effective visualisation. Since the survivorship rates of the heat stressed corals is so low, it might be worthwhile to have ambient and heat stressed survivorship plots on separate plots so the heat stress plot can have a more substantial y-axis. | This has been made clearer in the figure legend. We respectfully elect to have the values together to aid in the comparison of scale. | Lines 951-954:    “Median survivorship data is presented in Figures 2 and 4D, whereas survival data here is presented as the number of individual colonies per percent survival value. Survival at ambient and hot are shown together to aid in comparison.” |
| 2.25 | Figure 6D-F | Include space after reef name in vertical facet | The space has been added. | See New Figure 6 |
